# Improved Interpretability of Brain-Behavior CCA with Domain-driven Dimension Reduction

**DOI:** 10.1101/2021.07.12.451975

**Authors:** Zhangdaihong Liu, Kirstie J. Whitaker, Stephen M. Smith, Thomas E. Nichols

## Abstract

Canonical Correlation Analysis (CCA) has been widely applied to study correlations between neuroimaging data and behavioral data. Practical use of CCA typically requires dimensionality reduction with, for example, Principal Components Analysis (PCA), however, this can result in CCA results that are difficult to interpret. In this paper, we introduce a Domain-driven Dimension Reduction (DDR) method, reducing the dimensionality of the original datasets combining human knowledge of the structure of the variables studied. We apply the method to the Human Connectome Project S1200 release and compare standard PCA across all variables with DDR applied to individual classes of variables, finding that DDR-CCA results are more stable and interpretable, allowing the contribution of each class of variable to be better understood. By carefully designing the analysis pipeline and cross-validating the results, we offer more insights on the interpretation of CCA applied to brain-behaviour data.

## 1. Introduction

Complex, large-scale health projects such as the Human Connectome Project (HCP) (Van Essen et al. [2013]) and UK Biobank (Sudlow et al. [2015]) collect health related data from cohorts representative of a broad population that make the neuroimaging data even more valuable.

With such data, a central goal is to understand the interplay between the brain imaging and non-brain imaging variables. Canonical Correlation Analysis (CCA) (Hotelling [1936], Thompson [2005]) is a widely used tool to study such relationships. Smith et al. [2015] and Kumar et al. [2017] applied CCA to study the correlation between behavioral/demographic measures and brain imaging (resting-state fMRI) using HCP and UK Biobank data respectively. CCA, and closely related technique PLS (Partial Least Squares), have been applied to many other studies to investigate the links between neuroimaging data and other modalities (Krishnan et al. [2011], Sui et al. [2010], Vidaurre et al. [2017], Friman et al. [2001], Grellmann et al. [2015] and Whitaker et al. [2016]).

CCA takes in two sets of data and discovers the optimal combination of each set of variables to maximise correlation. To reduce the impact of noise and to avoid a degenerate solution when the number of subjects is less than the number of variables, a dimension reduction is often applied to each dataset and the reduced data are fed into CCA. Smith et al. [2015] and Kumar et al. [2017] applied Principal Component Analysis (PCA) to both behavioral/demographic measures, also known as subject measures (SM) and resting-state fMRI (brain measures; BM) to reduce the dimensionality of both to 100. However, those works provided no objective standard on the selection of the dimension to reduce to, and dimension reduction introduces further interpretability issue with the CCA results.

In this paper, we propose an alternative method of dimension reduction: Domain-driven Dimension Reduction (DDR). The main idea is to divide the data into sub-domains by function, then reduce the dimension on each of the sub-domains of SM and BM. Each sub-domain may contain a different number of variables and require different levels of dimension reduction. To this end we apply a two-way Cross-Validation (CV) method which estimates the dimensionality automatically by minimizing Predicted Residual Error Sum of Squares (PRESS). We then apply CCA to the DDR reduced SM and BM to study the correlations between brain and behaviour. To further improve the interpretability, orthogonal factor rotation is applied during dimension reduction.

We apply this analysis pipeline to the HCP S1200 release with 1003 subjects. The performance is assessed by examining canonical correlations, significant canonical variables and canonical loadings (also known as the structural coefficients). DDR offers us insights on the structure of sub-domains of SM and BM, and more interpretable CCA results. We carefully describe and apply a CV framework to assess the stability of DDR and CCA, by applying 5-fold CV.

For comparison, and to test the stability of the results, we replicated the analysis pipeline used by Smith et al. [2015] (PCA followed by CCA) on the larger S1200 data, and then compared its results with DDR CCA.

## 2. Method

### 2.1. Data

We used *N* = 1003 subjects from the Human Connectome Project (HCP) S1200 release. For brain measures (BM), we used the connectivity matrix (partial correlation) generated from resting-state fMRI data. Details of data acquisition can be found at the HCP database website (http://humanconnectome.org/data) and in Smith et al. [2013]. For non-imaging data, we considered 234 behavioral and demographic measures, and refer to them as subject measures (SM).

#### BM pre-processing

To generate BM, we used the pre-processing on resting-state fMRI (rfMRI) as described in Smith et al. [2015], and the pre-processed BM is available for download. In brief, Group-ICA (Independent Component Analysis) was performed to parcellate the brain using a 200-dimensional ICA parcellation. Each subject’s rfMRI data was then regressed against this to obtain one time series per ICA region. A functional connectivity matrix for each subject was generated by calculating the Tikhonov-regularized (Tikhonov [1963]) partial correlation for every pair of the time series. This resulted in a 200 × 200 connectivity matrix for every subject, and each of the entries represents a connectivity edge between two ICA regions.

#### Sign-flipping to maximise SM alignment

To facilitate interpretation of the variable loadings produced by CCA, we flipped the signs of some SM variables to provide a consistent meaning, specifically so that more positive values corresponded to ‘better’ life measures/outcomes. We first selected a benchmark variable ‘income’ and flipped variables that have negative correlations with it; we then examined the definition of each variable, flipping variables that did not already correspond to positive life outcomes. Note that flipping the signs of variables does not change the magnitude of covariance nor the eigenvalues, therefore does not affect the dimension reduction and CCA (proofs are shown in Theorem 2 and 3 in Appendix B.1). See Appendix B.2 for correlation matrices before and after sign-flipping.

#### Quality control and de-confounding

We removed ill-conditioned SM variables according to three criteria: if they had more than 50% missing values; if the standard deviation was 0; if more than 95% of the total entries were identical values. This left us 234 SM variables (see Appendix A for a full list of SM variables after quality control).

Both datasets were normalized by rank-based inverse normal (Blom [1958]) transformation (Beasley et al. [2009]) and then de-confounded. 15 confounding variables were carefully chosen as those that could potentially affect the relationship between brain and behaviour, including age, gender, height, weight and rfMRI head movement; and squared values for some of these variables such as age and BMI (see Appendix C for the full list of confounders). De-confounding was applied identically to each set of imaging and non-imaging variables, obtained by the residuals from a linear regression on the confound variables.

#### Grouping of SM and BM into sub-domains

All 234 SM variables were grouped into 14 sub-domains: Alcohol Use, Alertness, Psychiatric History, Tobacco Use, Drug Use, Emotion, Cognition, Family History, Physical Health, Motor, Personality, Sensory, Female Health and Demographics (including SES); this grouping followed the official HCP variable dictionary (https://wiki.humanconnectome.org/display/PublicData/HCP+Data+Dictionary+Public-+Updated+for+the+1200+Subject+Release). BM was grouped based on the 200 different ICA regions mentioned above. Thus, there are 200 BM sub-domains and each contain 200 brain edges. Each brain edge appears twice, one per (two) linked ICA regions, but this redundancy is accounted for in the dimension reduction.

### 2.2. Domain-driven Dimension Reduction

Domain-driven Dimension Reduction (DDR) is a refined application of Principal Component Analysis (PCA). Instead of reducing the dimension on the whole data space, SM and BM were grouped as above. PCA was then applied to each sub-domain in turn. The Principal Components (PCs) from the sub-domain analysis were concatenated to form the dimension-reduced data space. The dimensionality of the sub-domains are automatically estimated by minimizing the Predicted Residual Error Sum of Squares (PRESS) using a two-way Cross-Validation (CV) method (Bro et al. [2008]). An overview of the method is shown in Fig.1, and the illustration of the two-way CV is shown in Fig.2.

**Figure 1.**
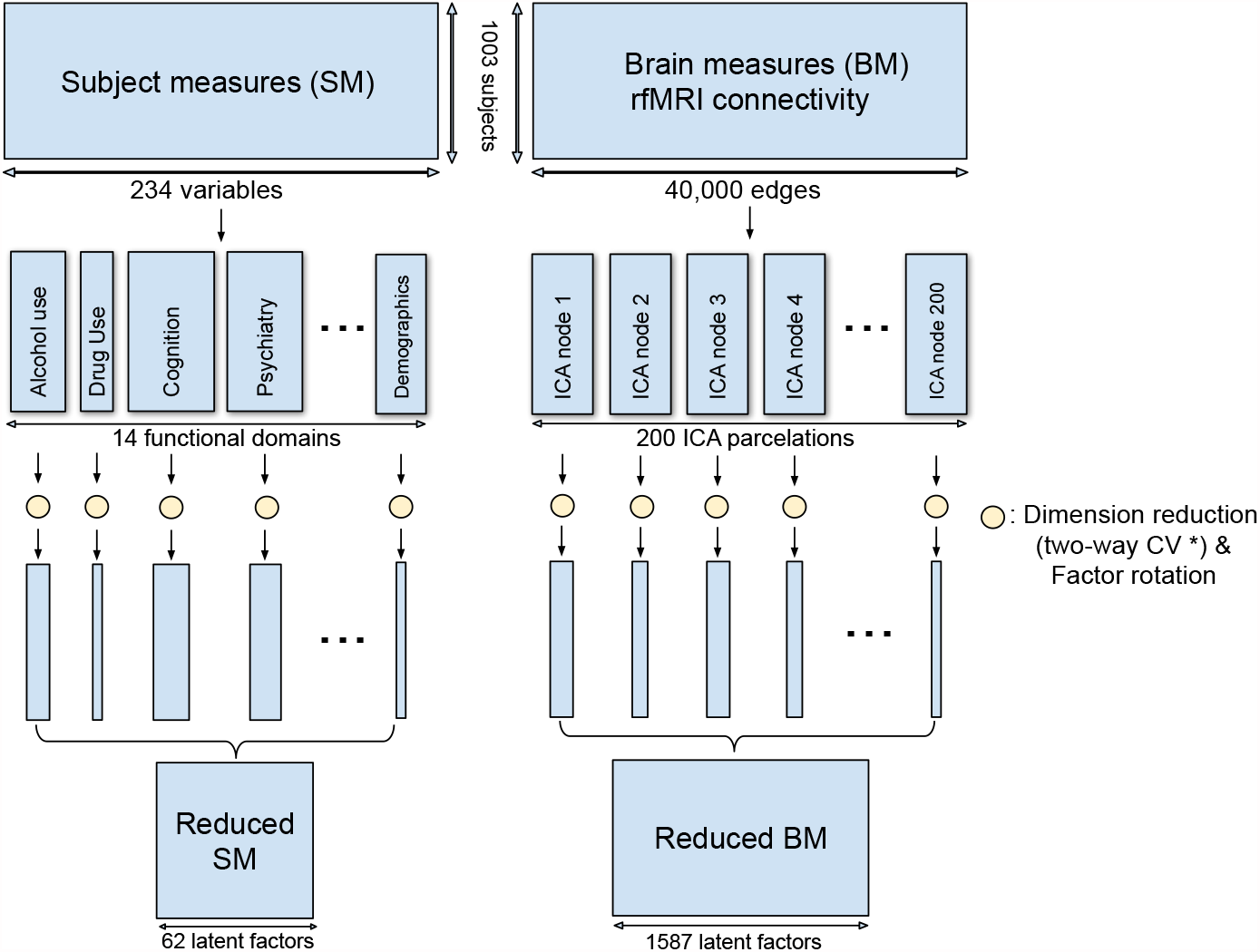
Method overview of DDR. SM and BM are first grouped into sub-domains. PCA is applied to each sub-domain while a two-way CV method (* see Fig. 2) is used to estimate the dimension. The rotated principal components from all sub-domains are concatenated to form the reduced SM and BM.

**Figure 2.**
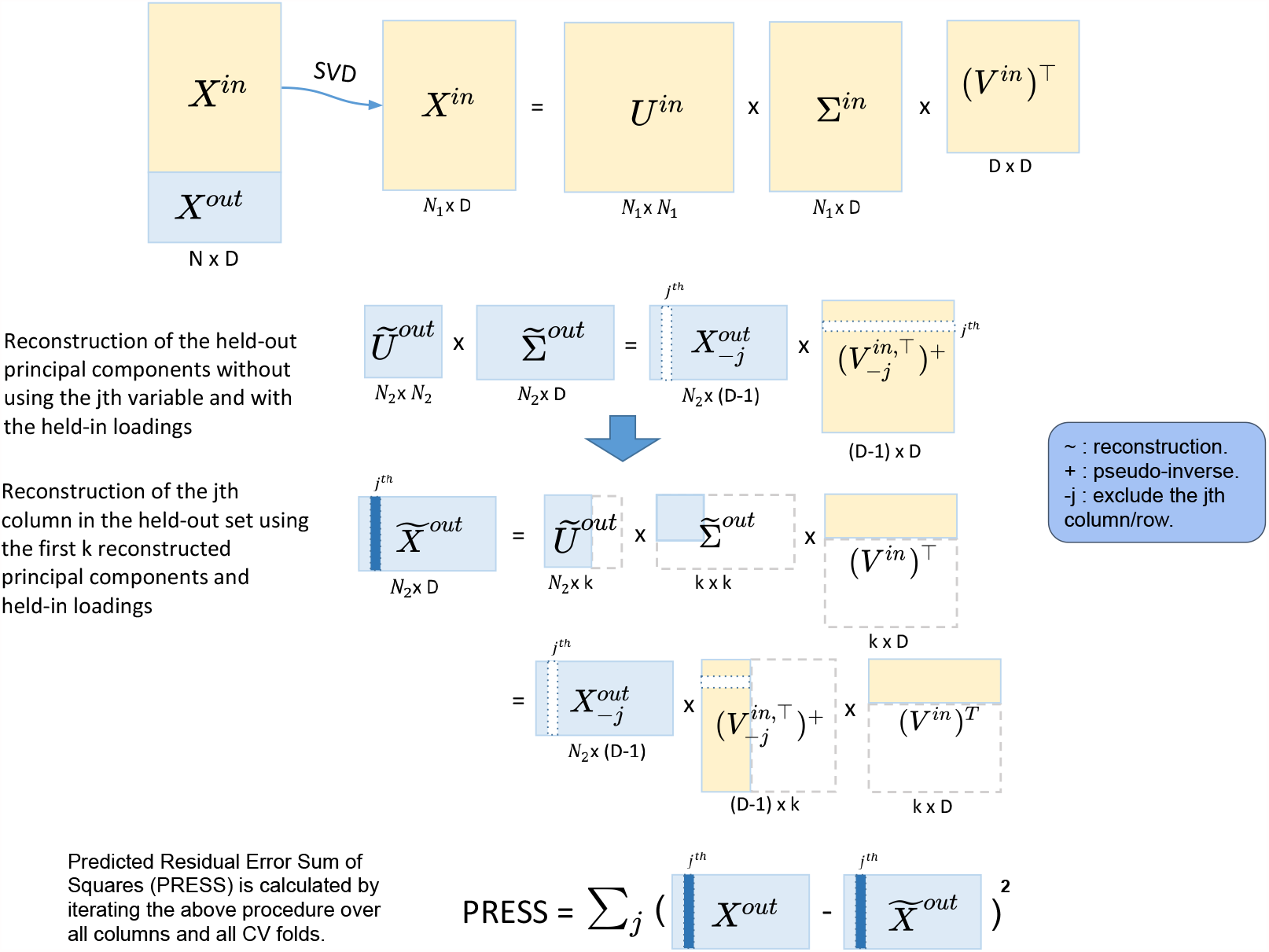
Illustration of the 5-fold two-way cross-validation. It minimizes PRESS and estimates the dimensionality in an automated fashion. Yellow blocks represent the training data and light blue blocks represent the test data. Two-way CV includes a subject-way (CV over subject direction) and a variable-way (CV over variable direction). Prediction error is calculated by the reconstruction error using different numbers of principal components.

#### Two-way cross-validation (CV)

The CV for dimension reduction is conducted variable-wise and subject-wise: a 5-fold subject-wise CV is performed, and within each of the 5 folds, a leave-one-out variable-wise CV is implemented. Dimension estimation is achieved by calculating PRESS for each of the held-out predictions (predicting with different dimensionalities, i.e. 1, 2, 3… PC(s)), selecting the dimension with the lowest PRESS as optimal.

In more detail, for the subject-wise 5-fold CV, we split the data into the held-in set (4/5 of the cohort), *X*^*in*^ and the held-out set (1/5 of the cohort), *X*^*out*^. Let *P* be the number of total variables in the sub-domain. Then we apply Singular Value Decomposition (SVD) to *X*^*in*^,

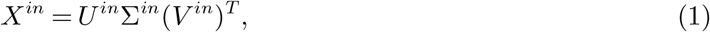

where *U*^*in*^ is the left-eigenvector matrix of *X*^*in*^, the eigenvectors of the subject covariance 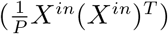;*V*^*in*^ is the right-eigenvector matrix, the eigenvectors of the variable covariance 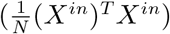; and Σ^*in*^ is the singular value matrix. The PCs of *X*^*in*^ are the singular-value-scaled left-eigenvectors, which we denote 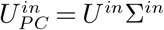, so that Eqn. 1 becomes

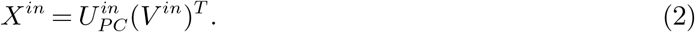

Noting that 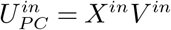 are the observations in the PC space. In order to reduce the dimensionality in the PC space to *k* (*k < P*), we can apply the following transformation

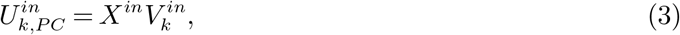

where 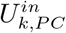 and 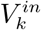 are the first *k* columns in 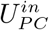 and *V*^*in*^, respectively.

Then we can likewise transform the held-out data to reconstruct the first *k* held-out PCs with the k-dimensional held-in principal loadings:

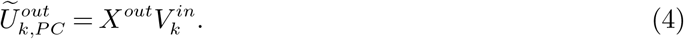

The lower dimensional reconstruction of the held-out data is thus

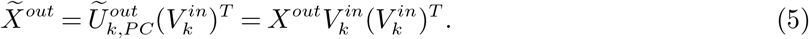

We can now calculate the prediction error as the difference between *X*^*out*^ and 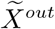. Iterating this algorithm over all 5 folds gives the subject-wise action of our two-way CV method, and we get a PRESS of

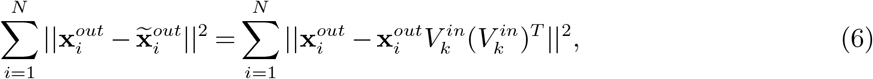

where **x**_*i*_ is a row vector and represents the *i*th subject in the held-out set.

However, the PRESS in Eqn. 6 monotonically decreases as *k* (the number of PC) increases and so is not suitable for dimensionality estimation. This is because the reconstruction of *X*^*out*^ in Eqn. 6 uses *X*^*out*^ itself. To address this we modify the reconstruction of 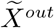 in Eqn. 5, predicting the *j*th column of *X*^*out*^ using the rest columns in *X*^*out*^:

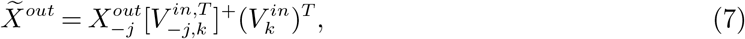

where 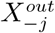 is *X*^*out*^ with the *j*th column removed; 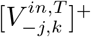 is the pseudo-inverse of the transpose of 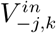, where 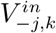 takes the first *k* columns of *V* ^*in*^ and then removes the *j*th row. The pseudo-inverse is required since removing a row of *V* ^*in*^ breaks its orthogonality. The *j*th column in 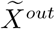 is now reconstructed without using the *j*th column in *X*^*out*^, and we denote this column as 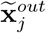. If we iterate *j* from 1 to *P*, we reconstruct the whole held-out set in turn. This is the variable-wise action in the two-way CV method. For each of the held-out in a CV fold, the corresponding PRESS can be calculated as:

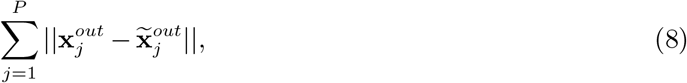

where 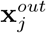 is the *j*th column in *X*^*out*^, and 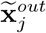 is as described above. Finally, the total PRESS for all subjects is calculated by summing PRESS in Eqn. 8 over all CV folds, completing the subject-wise action of the method.

Finding the dimensionality *k* with the minimum PRESS over dimensions completes the method for a given sub-domain. The reduced datasets for SM and BM are then obtained by the concatenation of the selected PCs from each of the sub-domains.

### 2.3. Evaluating the stability of DDR

Since DDR is based on CV, different random folds will give different PRESS values. Therefore, we repeated the two-way CV for DDR 50 times and took the mode of the estimated dimension for each sub-domain.

To further test the accuracy/rationality of the dimension DDR estimates, we compared the results from DDR with eigen-spectrum and null eigen-spectrum on each of the sub-domains. The eigen-spectrum provides information on the variance explained by each of the eigenvectors. The null eigen-spectrum is obtained by shuffling the row values for each column independently in the original matrix, and calculating the eigen-spectrum of the shuffled matrix. It shows the amount of ‘background noise’ exist in the dataset. When the null eigen-spectrum exceeds the eigen-spectrum, we can interpret this as the background noise taking over the information. If the estimated dimension from DDR falls near where the null eigen-spectrum crosses the eigen-spectrum, we have convergent evidence for the dimensionality estimation.

### 2.4. Canonical correlation analysis on brain imaging and behavioral data

Canonical Correlation Analysis (CCA) is a multivariate statistical approach to infer the relationship between two sets of variables. It aims to construct latent factors that maximize the correlation between two sets of data, *X* and *Y*, with a common number of rows and possibly different numbers of columns. For column vectors *A* and *B*, CCA finds two sets of linear combinations *P* = *XA* and *Q* = *YB* that are maximally correlated with each other. *P* and *Q* are known as the canonical variables; *A* and *B* are the canonical weights for *X* and *Y* respectively (Borga [2001]). The correlation between *P* and *Q* is called the canonical correlation, *R*.

To evaluate the importance of variables, we use canonical loadings, also known as structural coefficients (Egloff et al. [2010], Borga [2001]), defined by correlating canonical variables with the observed datasets, in this case, SM (denoted as *X*) and BM (denoted as *Y*):

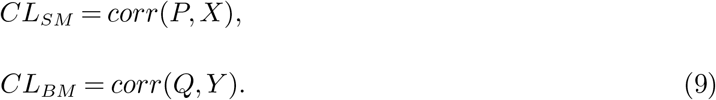

Permutation testing is used to test the significance of the canonical variables.

#### DDR CCA pipeline

Since we found that the DDR dimension for BM is still larger than the number of subjects and much larger than the DDR estimation for SM, we applied PCA to DDR reduced BM to further reduce its dimensionality. We reduced the dimension of BM to 100 to match the method in Smith et al. [2015], and also considered the same dimension as the DDR reduced SM.

To further improve the interpretation of these PCs, we applied Varimax factor rotation (Kaiser [1958]) to the principal loadings in the sub-domains. The rotated PCs, *RC*_*X*_ and *RC*_*Y*_, are then fed into CCA. Notably, orthogonal rotation is an invariant transformation on CCA inputs. Therefore, it does not affect CCA outputs, only improves the interpretation of the DDR factors.

We examined the number of significant pairs of canonical variables using permutation testing, and evaluated the variable importance by two different measures: Canonical loadings for observed variables, and canonical loadings for DDR factors (CCA inputs). For the first measure, we calculated the same loadings as in Eqn. 9. The second set of loadings offers insights in the importance of each sub-domain, and are calculated as:

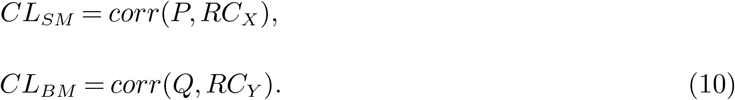

We have also calculated variance explained by each of the significant canonical variables in the original datasets of SM and BM. The R-squared value was computed for each variable and then averaged.

### 2.5. Stability study of CCA

In order to test the stability of the CCA results, we applied 5-fold CV (Fig. 3). For each fold, we tested our model on the training set, four fifths of the data (∼800 subjects), and validated on the test set, one fifth of the data (∼200 subjects). The splits do not break the families, i.e. subjects from the same family will go into the same group. The detailed procedure of CV is shown in Algorithm 1.

**Figure 3.**
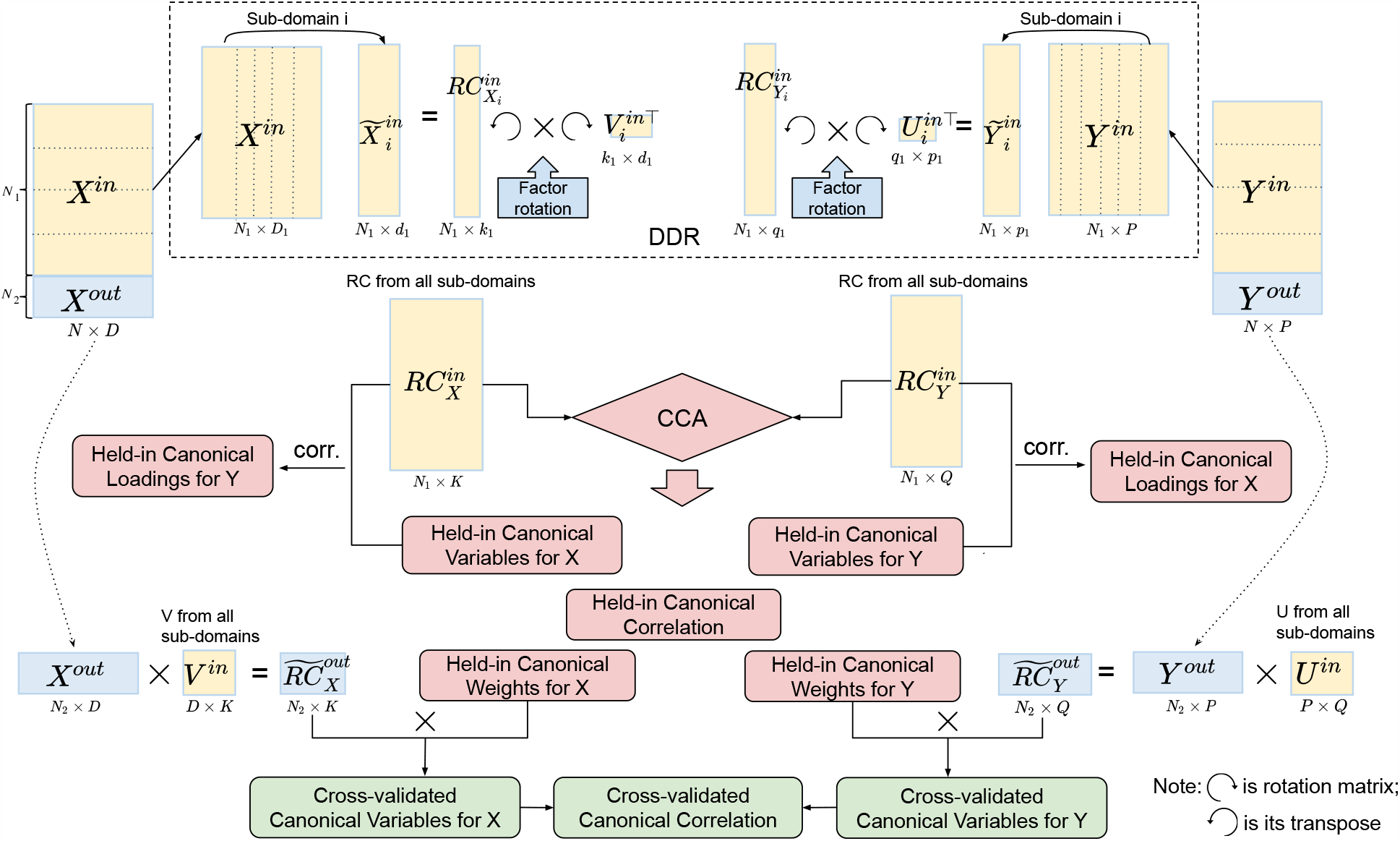
Illustration of 5-fold cross-validation (CV) in the DDR CCA analysis. We apply DDR (dotted box on the top) to the training set after the split of the data. Within DDR the principal loadings are rotated to construct the rotated components (RCs), and the RCs from the training set are fed into CCA. Cross-validated canonical variables and correlations are obtained by multiplying the training canonical weights with the RCs from the test set.

### 2.6. Comparison between PCA and DDR

To assess the performance of DDR in comparison with PCA, we applied the same analysis pipeline using PCA instead of DDR on the same datasets. Smith et al. [2015] applied PCA to reduce the dimensions of SM and BM to both 100. However, our DDR method automatically reduces the dimension of SM to under 100, and of BM to over 100. Trying to make the dimensions consistent, we apply PCA to match the dimension of DDR reduced SM dataset. For BM, we chose to apply PCA after DDR to reduce BM dataset to 100.

We compared the variance explained by the PCs obtained by PCA and DDR respectively in the original SM and BM spaces. Feeding PCA reduced datasets and DDR reduced datasets into CCA separately to compare their canonical variables, canonical correlations and canonical loadings, as well as the variance explained by canonical variables in the original SM and BM spaces. We also applied the same CV procedure (as shown in Algorithm 1 with the replacement of PCA with DDR) to compare the stability of PCA CCA with DDR CCA.

## 3. Results

### 3.1. PCA-based CCA

Tab. 1 is a summary table showing variance explained by significant canonical variables of SM and BM, canonical correlations and the number of significant CCA modes at 5 sets of different input dimensions. We notice that the first canonical variable does not necessarily explain the most variance in the observed datasets. Interestingly, Tab. 1 shows that if we decrease the dimension(s) of SM and/or BM, the canonical variables would explain more variance in the observed dataset(s). For example, by observing the last three rows in Tab. 1, the input dimension of BM decreases from 100 to 30. The ‘Variance Explained (%) by BM Canonical Variable’ increases for all significant canonical variables. However the strength of canonical correlation decreases as the dimension decreases.

**Table 1.**
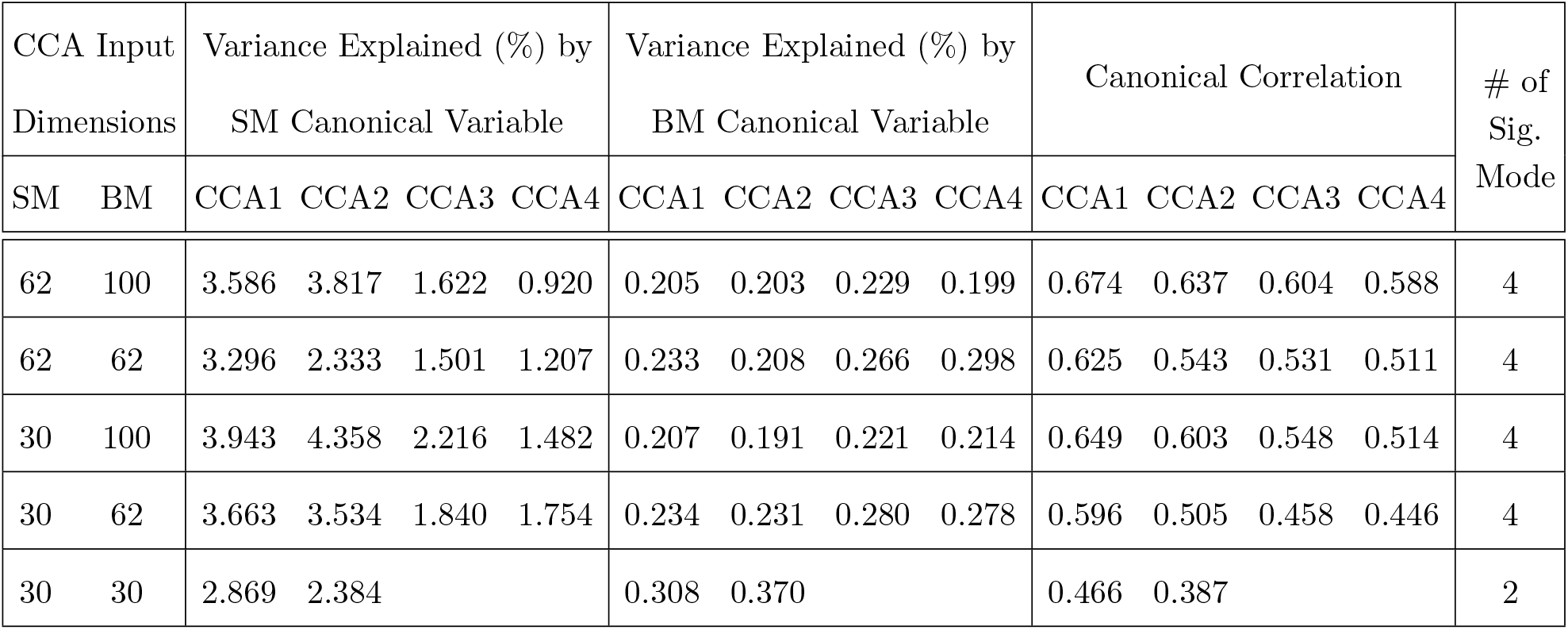
Summary table for PCA CCA with 5 different input dimensions of CCA (first column). Second and third columns show the variance explained by the SM and BM canonical variables in the observed SM and BM datasets for significant canonical pairs respectively; the fourth column shows the canonical correlation for the canonical pairs; the last column shows the number of significant canonical pairs obtained by permutation testing.

| CCA Input Dimensions |  | Variance Explained (%) by SM Canonical Variable |  |  |  | Variance Explained (%) by BM Canonical Variable |  |  |  | Canonical Correlation |  |  |  | # of Sig. Mode |
| --- | --- | --- | --- | --- | --- | --- | --- | --- | --- | --- | --- | --- | --- | --- |
| SM | BM | CCA1 | CCA2 | CCA3 | CCA4 | CCA1 | CCA2 | CCA3 | CCA4 | CCA1 | CCA2 | CCA3 | CCA4 |  |
| 62 | 100 | 3.586 | 3.817 | 1.622 | 0.920 | 0.205 | 0.203 | 0.229 | 0.199 | 0.674 | 0.637 | 0.604 | 0.588 | 4 |
| 62 | 62 | 3.296 | 2.333 | 1.501 | 1.207 | 0.233 | 0.208 | 0.266 | 0.298 | 0.625 | 0.543 | 0.531 | 0.511 | 4 |
| 30 | 100 | 3.943 | 4.358 | 2.216 | 1.482 | 0.207 | 0.191 | 0.221 | 0.214 | 0.649 | 0.603 | 0.548 | 0.514 | 4 |
| 30 | 62 | 3.663 | 3.534 | 1.840 | 1.754 | 0.234 | 0.231 | 0.280 | 0.278 | 0.596 | 0.505 | 0.458 | 0.446 | 4 |
| 30 | 30 | 2.869 | 2.384 |  |  | 0.308 | 0.370 |  |  | 0.466 | 0.387 |  |  | 2 |

#### Algorithm 1

Cross-validation procedure for the DDR CCA analysis

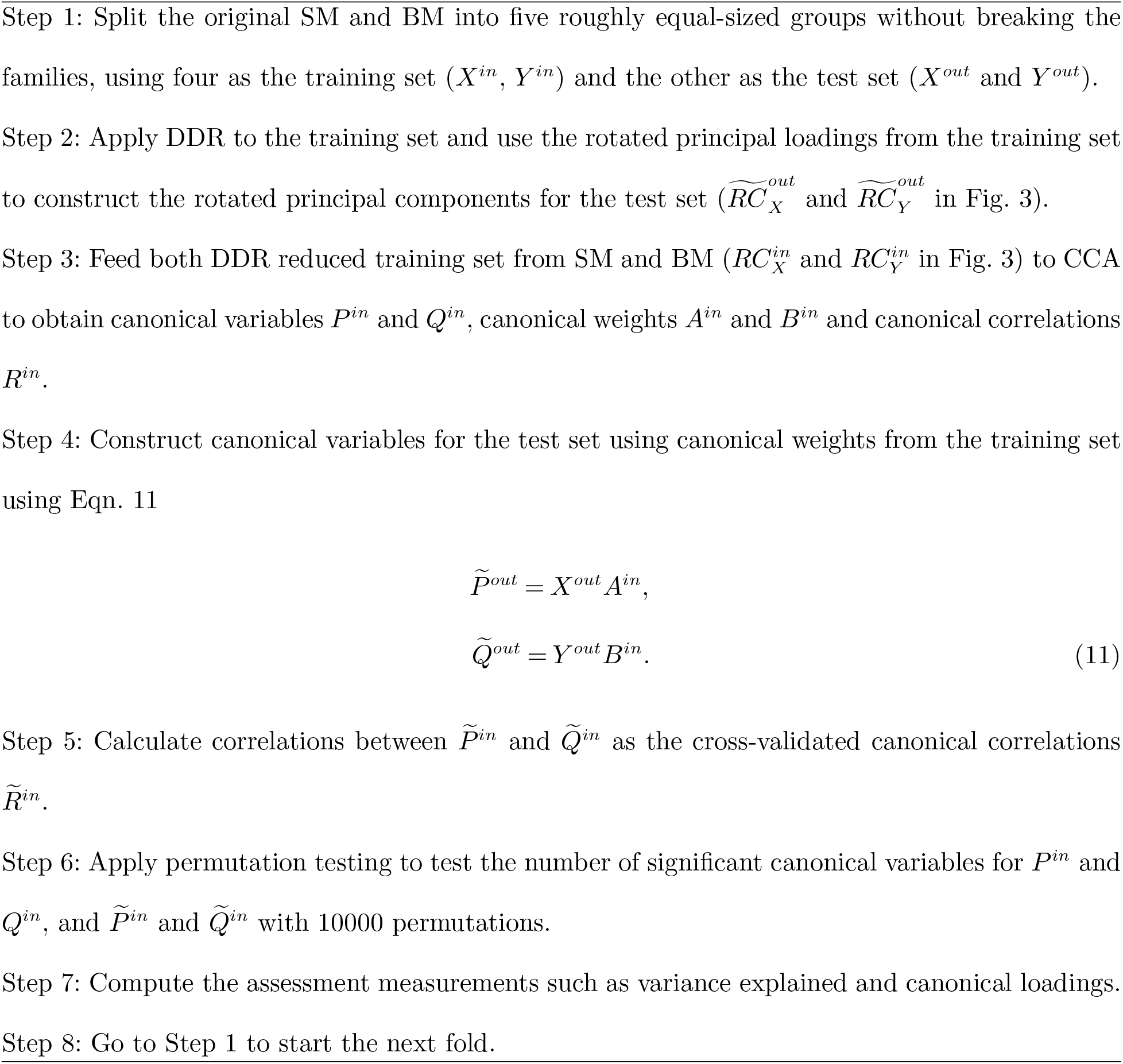

For the rest of the paper, we focus on the 62 dimensional SM and 100 dimensional BM since 62 is the DDR estimated dimension for SM and 100 was selected in previous studies (Smith et al. [2015]). There are 4 significant canonical pairs identified by permutation testing in this setting. The canonical loadings of SM (Fig. A.2) for these 4 canonical variables display 4 behavioral/demographic modes. The first set is mainly loaded on cognition variables; the second set is dominated by tobacco variables; most of the top loadings in the third set are alcohol variables; the fourth set is more of mixture with cognition, emotion and motor variables.

Note that most of the SM variable loadings shown in Fig. A.2 have the same sign, and this is in contrast to previous CCA results with the HCP data. For example, Smith et al. [2015] found a mode with tobacco use and education measures having opposing signs, while here, after flipping the signs of the observed variables, they are now on the same side of the axis (CCA mode 2 in Fig. A.2). While the canonical variable found by CCA is invariant to sign flips of the variables, the canonical loadings of course reflect any sign flips (see Theorem 3 in Appendix B.1).

### 3.2. DDR results

We generated a summary report for each of the 14 sub-domains (Appendix E) to helps us understand the structure of each sub-domain.

Two of the panels in the Family History report are shown in Fig. 4 to show as an example. Top subplot in Fig. 4 shows the rotated principal loadings, i.e. the variable importance on generating the latent factors for the sub-domain. We observed higher interpretability on the rotated loadings, therefore, used them to summarise the meaning of latent factors in the sub-domain, as shown in Tab. 3. The dimension of the sub-domains is decided by the minima of the red line in the bottom subplot in Fig. 4, which is calculated by Eqn. 8. These DDR estimations also generally corresponded to where the actual and null eigenspectrum cross (Panel A in Fig. A.3 to A.16 in Appendix E).

**Figure 4.**
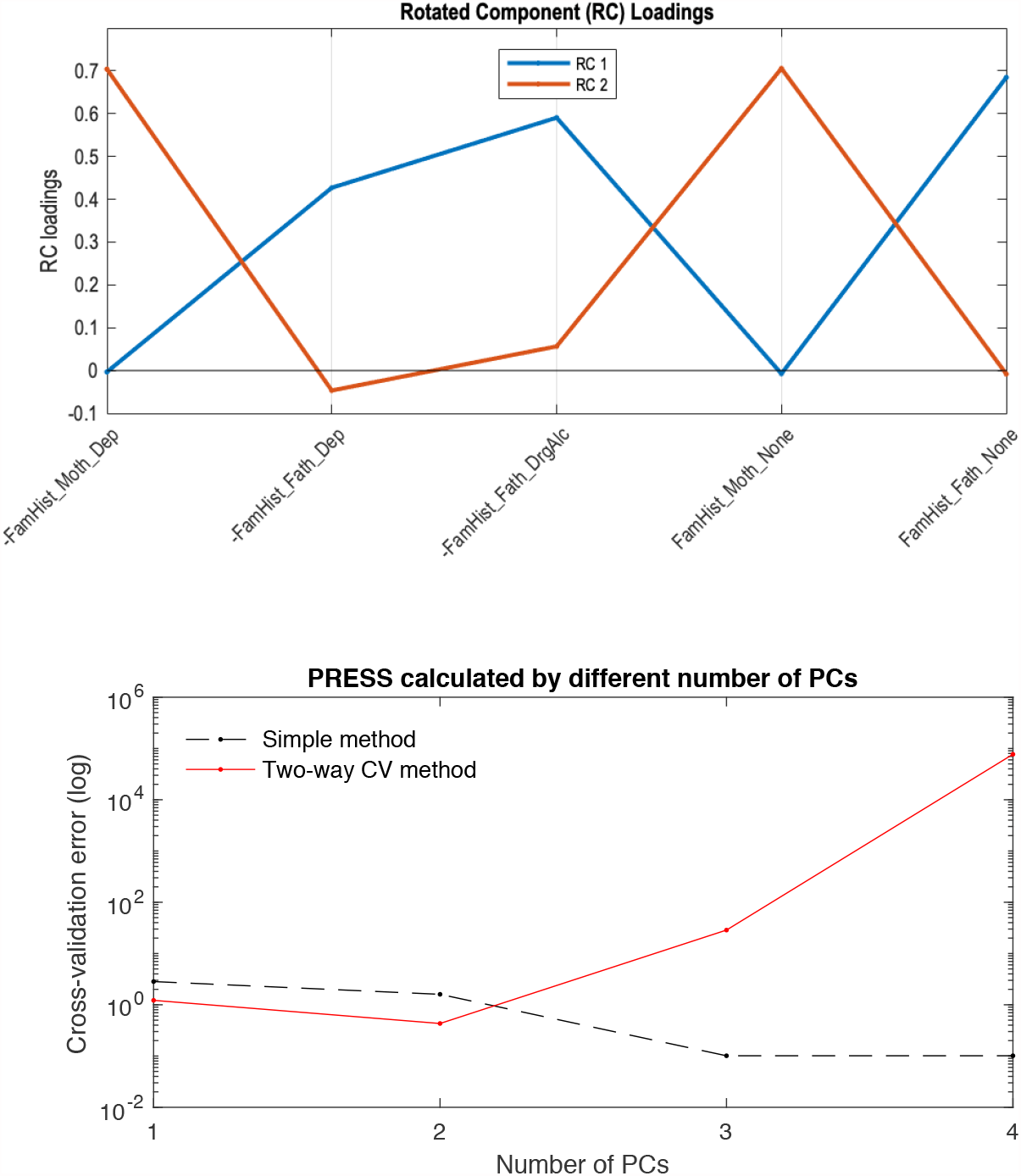
Top figure shows the rotated principal loadings; figure at the bottom shows the error curves calculated by Eqn.6 (dotted line) and Eqn.8 (red line), with the minimal error circled at the second component. The naive way of calculating PRESS (dotted line) is monotonically decreasing, while the two-way CV method (red line) offers a minimum point.

Moreover, by investigating the sub-domain structures, we observed strong stability of DDR factors and understood better the composition of the latent factors in each sub-domain.

In total, DDR selected 62 SM factors from the 14 sub-domains. Different from choosing PCs by a simple cut-off point from the variance explained point of view, we can see factors in different domains explain different amount of variances (third column, Tab. 2). For example, the first PC (out of 10) in Tobacco Use explains 80.54% variance in the whole sub-domain, and the first PC (out of 11) in Drug Use explains less than 50%. However, with only 1 PC in these two sub-domains, they achieved the lowest prediction errors in the test set. For BM, DDR reduced 200 ICA regions/sub-domains from total dimension of 40000 to 1587.

**Table 2.**
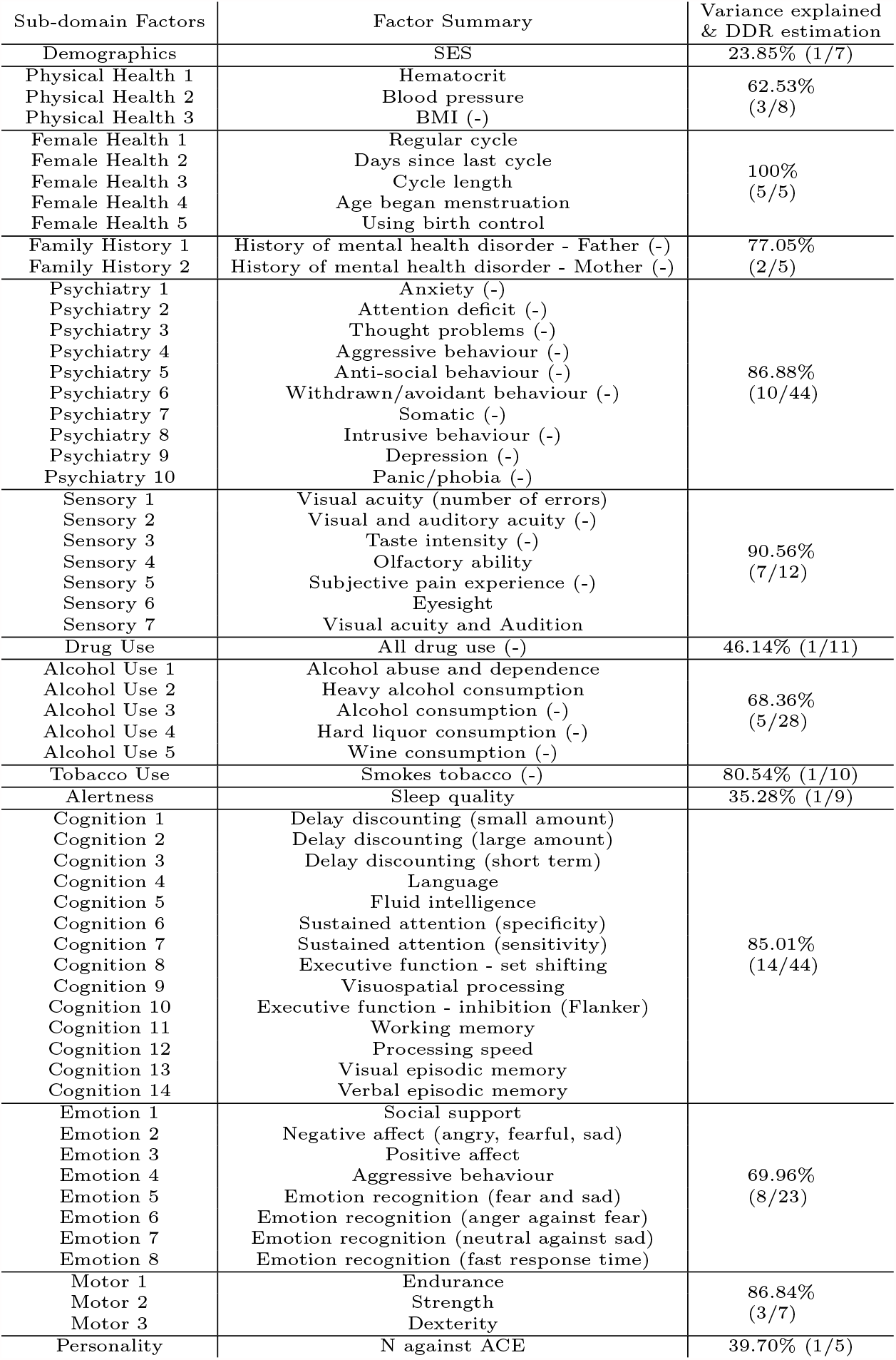
Summary of SM sub-domains. The factors are orthogonally rotated principal components and ordered by R-squared values in the original sub-domain. Second column shows the factor names summarized from panel E in each of the sub-domain report like Fig. A.3. The third column shows the variance explained by the dimension reduced sub-domain in the original sub-domain. Numbers in brackets are the two-way CV estimated dimension verses the total number of variables in the sub-domain. Personality is summarized by Big Five personality traits: Neuroticism (N), Agreeableness (A), Extraversion (E), Conscientiousness (C), Openness to experience (O).

**Table 3.** Summary table for DDR CCA. The first column shows the dimensions of SM and BM as CCA inputs; second and third columns represent the variance explained by the SM and BM canonical variables in the observed SM and BM set for significant canonical pairs respectively; the fourth column shows the canonical correlation for the canonical pairs; the last column shows the number of significant canonical pairs.

| CCA Input Dimensions |  | Variance Explained by SM Canonical Variable |  |  | Variance Explained by BM Canonical Variable |  |  | Canonical Correlation |  |  | # of Sig. Mode |
| --- | --- | --- | --- | --- | --- | --- | --- | --- | --- | --- | --- |
| SM | BM | CCA1 | CCA2 | CCA3 | CCA1 | CCA2 | CCA3 | CCA1 | CCA2 | CCA3 |  |
| 62 | 100 | 2.629 | 1.727 | 1.698 | 0.202 | 0.183 | 0.211 | 0.632 | 0.582 | 0.574 | 3 |
| 62 | 62 | 2.318 | 1.386 | 2.361 | 0.203 | 0.302 | 0.271 | 0.555 | 0.519 | 0.495 | 3 |
| 30 | 100 | 3.323 | 3.928 | 1.737 | 0.194 | 0.181 | 0.267 | 0.586 | 0.541 | 0.503 | 3 |
| 30 | 62 | 2.776 | 2.867 | 1.290 | 0.306 | 0.260 | 0.244 | 0.475 | 0.445 | 0.440 | 3 |
| 30 | 30 | 2.716 |  |  | 0.376 |  |  | 0.419 |  |  | 1 |

### 3.3. DDR-based CCA

Similar to the PCA-based CCA analysis, we applied DDR-based CCA analysis to different input dimensions of CCA (Tab. 3). Same rule in PCA-based CCA was found here: lower CCA input dimension leads to canonical variables explaining more variance, whereas the canonical correlations get weaker. Comparing each setting with results in PCA-based CCA (Tab. 1), we found that the number of significant canonical variables is always one lower than the PCA case.

**Table 4.** 5-fold cross-validation on 62 dimensional SM and 100 dimensional BM in DDR CCA analysis. Mean variance explained and canonical correlations are shown for the first 3 pairs of canonical variables with standard deviation (std) in brackets.

|  | CCA 1 | CCA 2 | CCA 3 |
| --- | --- | --- | --- |
| Mean variance explained in held-in SM set (std) | 2.34% (0.50) | 2.13% (0.85) | 1.95% (0.84) |
| Mean variance explained in held-in BM set (std) | 0.23% (0.027) | 0.22% (0.028) | 0.22% (0.017) |
| Mean variance explained in CV SM set (std) | 2.80% (0.73) | 2.60% (0.85) | 2.37% (0.57) |
| Mean variance explained in CV BM set (std) | 0.63% (0.051) | 0.65% (0.086) | 0.65% (0.059) |
| Mean canonical correlation for held-in set (std) | 0.662 (0.011) | 0.639 (0.016) | 0.611 (0.0084) |
| Mean canonical correlation for CV set (std) | 0.228 (0.037) | 0.108 (0.057) | 0.174 (0.039) |

#### Canonical loadings for SM

For 62-dimensional DDR SM and 100-dimensional DDR+PCA BM, permutation testing identified three significant canonical pairs. The top 20 canonical loadings (in absolute value) for each of them are shown in Fig. 5. Noticeably, all top 20 loadings for these three canonical variables are positive after sign-flipping of the observed variables.

**Figure 5.**
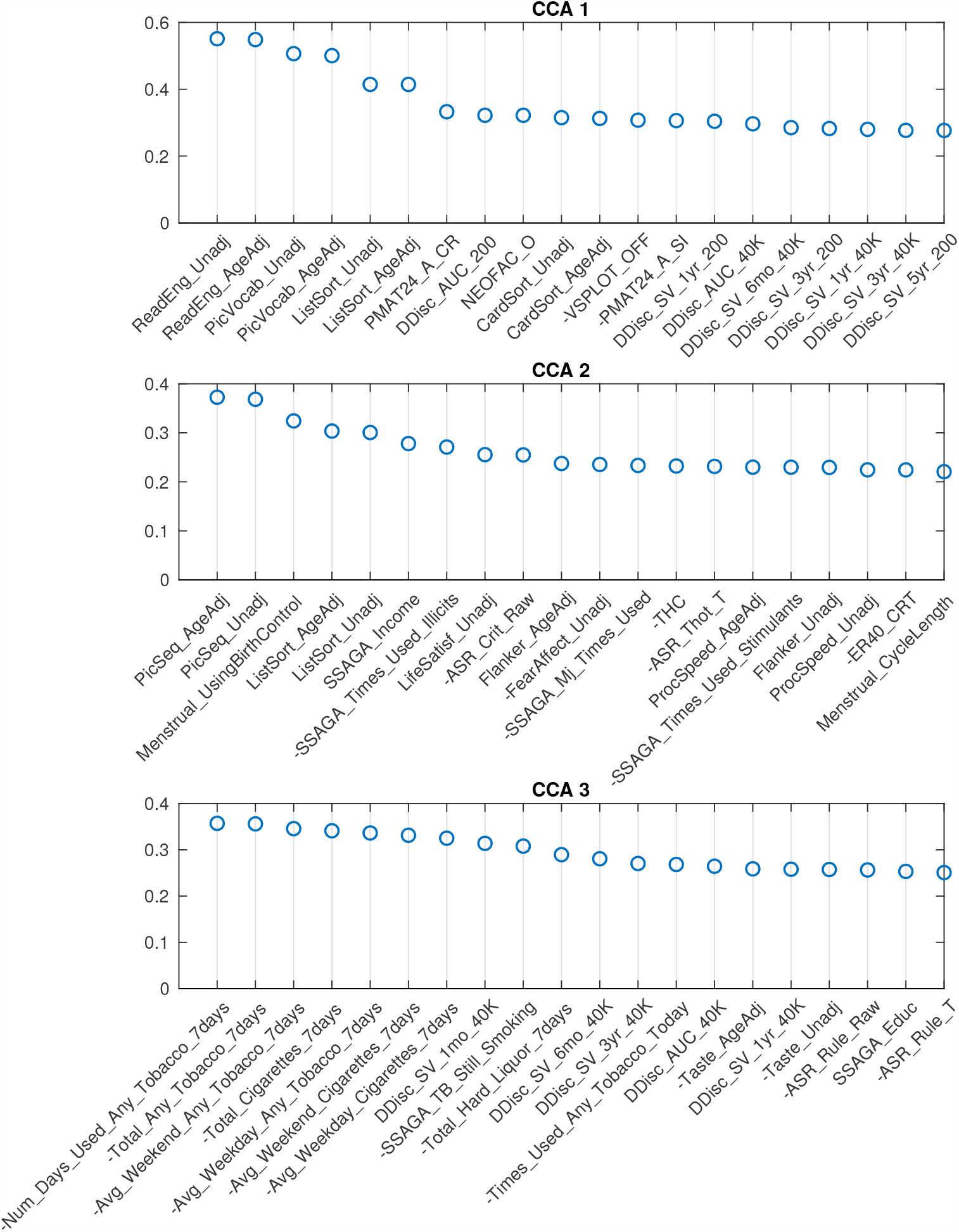
Top 20 SM canonical loadings for 3 significant canonical variables. Variable name with ‘-’ sign shows that it was flipped in the original dataset. Canonical loadings of CCA 1 are very similar to the first set of PCA CCA, heavily cognition dominated; the second set is mixed with cognition, drug use etc; The third set is combination of tobacco use and cognition variables.

With the help of DDR, we are able to explore the contributions of CCA inputs directly, by calculating the canonical loadings of them using Eqn. 10. The canonical loadings of the inputs are not interpretable in PCA-based CCA. However with DDR, we are able to interpret not only the latent factors but also the canonical loadings on those factors (Fig. 6). Using the summarized latent factors in Tab. 2, we are able to conclude, for example, in the first set of canonical loadings (the first subplot in Fig. 6), Language factor (Cognition 4) has the largest loading. The second and third largest loadings are Cognition 3 and 1, and they are Delay Discounting factors.

**Figure 6.**
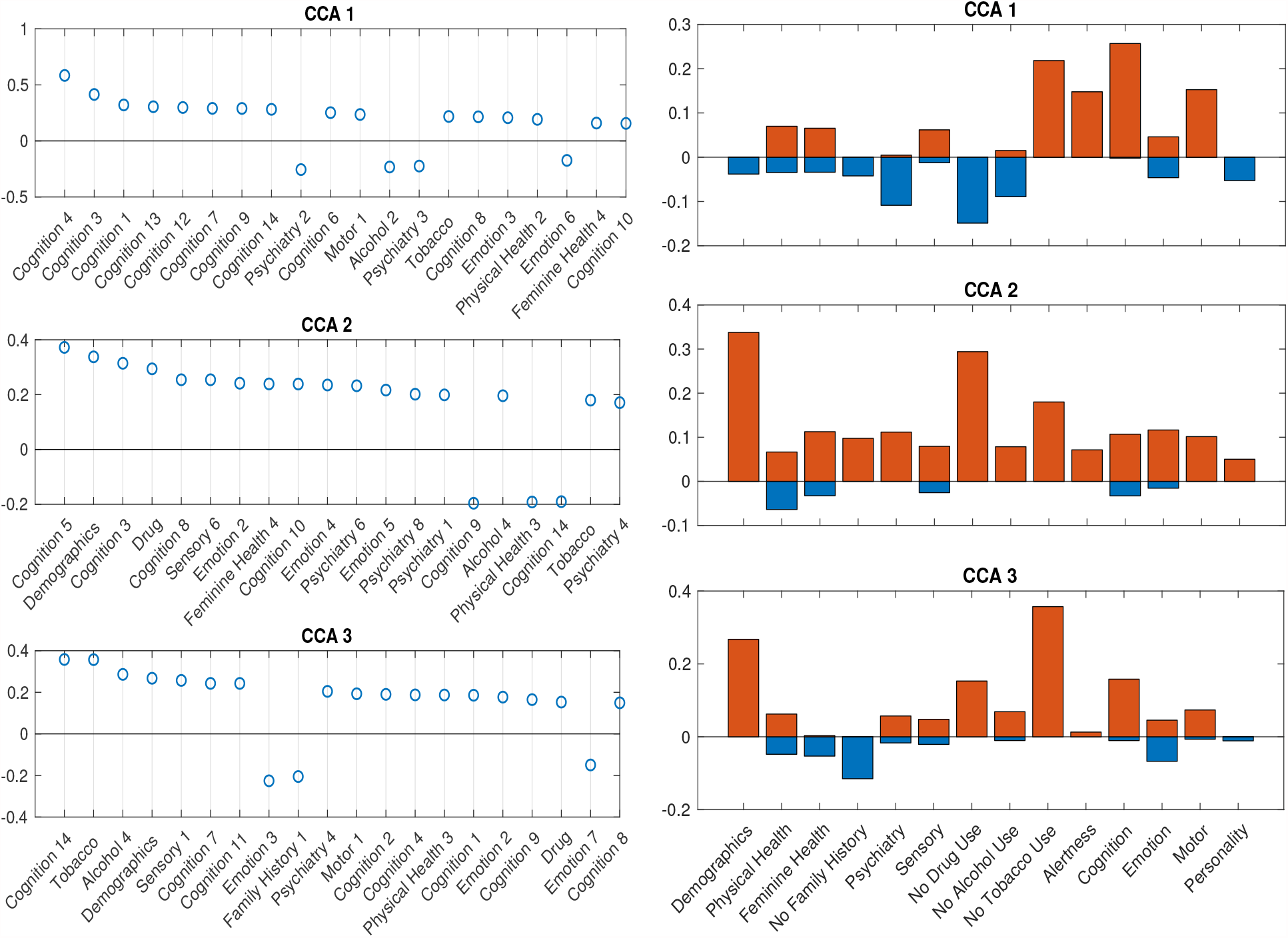
SM canonical loadings on the CCA input for 3 significant canonical variables. Left set of figures shows top 20 loadings for 3 significant canonical variables respectively; right set of figures show the mean of all positive loadings (red bars) and the mean of all negative loadings (blue bars) within each sub-domain for the 3 significant canonical variables.

The right set of figures in Fig. 6 offer us insight in the overall contribution of each sub-domain, by sign of their contribution: the total length of each blue-red bar pair is the average R-squared explained, with the contribution from positively-weighted variables plotted above the x-axis, negatively-weighted plotted below. We notice here the top loadings and overall loadings are not mono-signed anymore even with the sign-flipping in effect. Interestingly, the pattern presented in the first overall canonical loadings (top right, Fig. 6) is driven by people with good cognition and motor ability, who don’t smoke, but take drugs, have some kind of mental disorder and drink. The second and third sets are displaying good well-being patterns. In particular, the second set of loadings are dominated by high SES and no drug use; the third set shows the alignment between no tobacco use and high SES. All of these patterns cannot be observed by using PCA after sign-flipping.

#### Canonical loadings for BM

Each set of canonical loadings for BM is a 200×200 symmetric matrix. Each entry represents a CCA connection (edge) between two ICA regions. We first map this loading matrix with the signs of the group mean correlations between the ICA regions, i.e. if two ICA regions were negatively correlated at resting-state, it would decrease the positive CCA strength but enhance the negative CCA strength. Due to the difficulty of interpreting each of these 19,900 (200 ∗ 199*/*2) edges, we came up with the following summary statistics. Averaging the top 20 (10%) positive and negative modulated canonical loadings for each ICA region (in each column/row) as the positive and negative CCA strength respectively. They are shown in Fig. 7.

**Figure 7.**
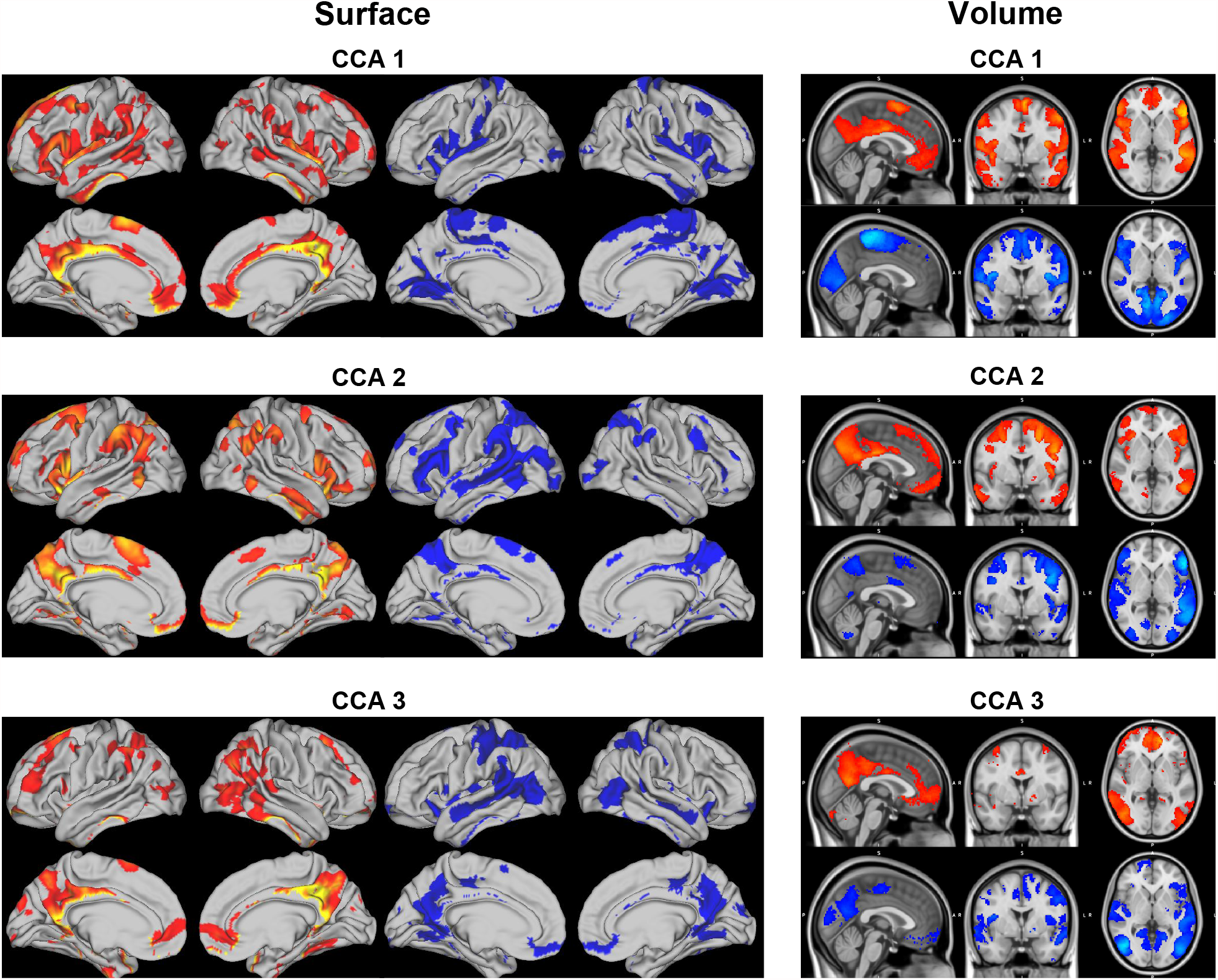
Positive and negative CCA strengths on brain surface (left) and volume (right) for the 3 significant canonical variables. The visualization is cut by 80 percentile. Positive (red maps) and negative (blue maps) CCA strengths are generated by mapping the canonical loading with the sign of population mean correlation between each pair of ICA regions, then average the top 20 positive and negative modulated loadings respectively.

The CCA strengths for CCA 1 ^1^ illustrate a weak contrast between language, sentences, semantic areas (positive strength) against premotor, motor, primary areas (negative strength); positive and negative strengths for CCA 2 ^2^ are much less distinguishable, both overlapping with parietal and intraparietal which are arguably linked to working memory and default mode network. The positive CCA strength maps for CCA 3 ^3^ overlap considerably with CCA 2. The positive map shops weak connection with the default mode network whereas the negative map activates in occipital and pre-motor areas.

Combining results from both SM and BM sides, CCA mode 1 reveals an interesting pattern: language and comprehension related brain areas associate positively with no tobacco use, no psychiatric illnesses, better alertness and cognitive ability, and negatively with drug use. Whereas drug use is positively correlated with the motor areas in the brain.

### 3.4. Stability of DDR-based CCA

We applied 5-fold CV to DDR CCA with 62-dimensional DDR SM and 100-dimensional DDR+PCA BM. We also ran permutation testing on the training and cross-validated sets for 10000 simulations to get significant canonical pairs. Permutation testing resulted in mostly 2 significant canonical pairs on the training sets around, and ranged from 0 to 4 on the CV sets, with 0 or 1 being the common numbers.

In general, for canonical correlation, as the sample size gets larger, the correlation gets weaker. The first canonical correlation is 0.632 for 1003 subjects, and 0.662 on average for four fifths of those subjects. This also applies to the number of significant CCA pairs permutation test detects. With the HCP 500 release, only 1 significant pair was detected (Smith et al. [2015]); we replicated the study with HCP 900 release and found 2 pairs; In this study, 3 pairs were discovered using the whole cohort. However in CV, the training set consists four fifth of the subjects (the same amount as in the 900 release), and 2 is most common significant number (with the permuted mean canonical correlation being around 0.6, and standard deviation around 0.01). which is consistent with the previous finding. Hence, we are going to examine further the results on the 2 significant pairs on training sets in CV.

We have selected the top 20 SM canonical loadings in every fold and took the ones that occurred at least two times out of the 5 folds CV for the 2 significant canonical variables. Stability of the first and second sets of SM canonical loadings on the observed SM data is shown in Fig. A.17. Language variables turn to be the most stable and heavily weighted in the first set, appearing on the top in every fold, and most of the variables in the first set in CV appeared in the first canonical loadings for the whole cohort. The second set of canonical loadings turn to be less stable than the first one with the most occurrence being 3 out of 5. The second set of canonical loadings in Fig. A.17 look like a combination of the second and third sets in the whole-cohort analysis.

We focus on the stability of SM canonical loadings on the CCA input (Fig. 8). The top two most stable latent factors are Language factor (Cognition 4) and Delay Discounting (Cognition 3), and they are the top two in the one-off analysis (Fig. 6). Similar to the canonical loadings on the observed data, the second loadings here are combined from the second and third loadings in the whole-cohort analysis. Comparing stability of the canonical loadings on CCA input (Fig. 6) with the canonical loadings on observed variables (Fig. A.17), there is improvement from the occurrence frequency point of view, as well as in the variance of the canonical loadings across different folds. Particularly for the second set of canonical loadings, loadings on the observed variables have the highest occurrence being 3 whereas 5 on the CCA input. The only factor DDR chose in Demographics and Drug Use appeared in every single fold with high and stable canonical loadings, which we cannot observe from the loadings on the observed variables. Moreover, similar to the one-off analysis on the whole cohort, stability of canonical loadings on the CCA input presents the contrast of relationship again. Psychiatry, Drug, Alcohol factors have opposite contributions to Cognition and Motor ones in the first set of canonical loadings. (See Fig. A.18 for the stability on BM canonical loadings.)

**Figure 8.**
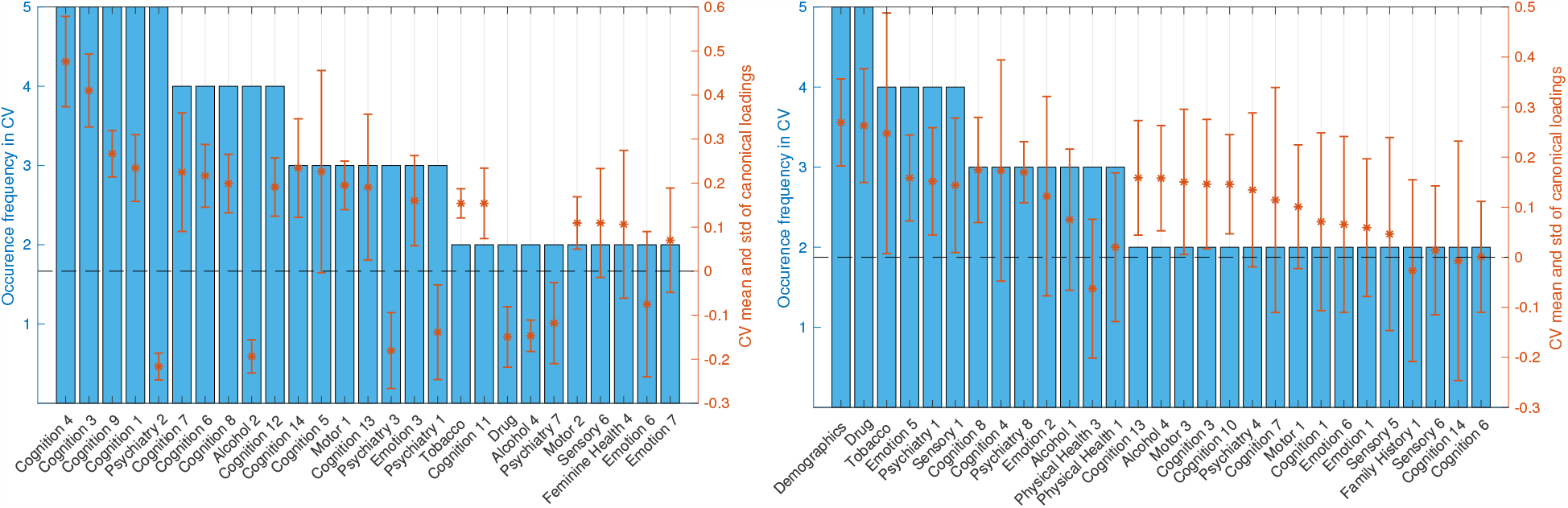
Stability of SM canonical loadings on CCA input. Bar plot shows the occurrence frequency in CV out of the 5 folds. Variables are chosen by selecting the top 20 mostly weighted ones in each fold. The ones appeared at least twice are shown above.right axis shows the mean and the standard deviation over all occurred loadings. Left and right plots are the canonical loadings for the first and second canonical variables respectively.

## 4. Discussion

In this paper, we carefully replicated the study in Smith et al. [2015] with a modified analysis pipeline for the HCP S1200 release. Comparing with the arbitrary choice of 100-dimension PCA in that work, we proposed a more automated way of estimating dimensionality of the data, particularly for the function-specific sub-domains of the SM and independent regions of BM. It is often quite challenging to interpret results of Canonical Correlation Analysis (CCA) applied to behavioral and brain imaging data, e.g. the canonical variables, canonical loadings, canonical correlations etc. The biggest motivation of proposing Domain-driven Dimension Reduction (DDR) is to improve the interpretibility of these results.

### Sign-flipping and de-confounding

Sign flipping all variables to align with positive life outcomes produced modes that were consistent with Smith et al. [2015], however, when plotted, the results appear different due to sign-flipping. In particular, the canonical loading of variables like the Picture Vocabulary Test, Oral Reading Recognition Test, Fluid Intelligence (correct response) now share the same sign as tobacco and alcohol measures (Fig. A.2 and Fig. 5).

Aside from sign-flipping, differences from previous HCP results may arise, in part, from the different variable sets used, as this work considered a wider range of variables. Other differences included slightly different set of confounders, with this analysis using racial factors, release versions, age and gender as additional confounders.

### Comparing DDR with PCA

By grouping the variables of SM into sub-domains based on their functions, we are able to interpret the canonical loadings of the inputs of CCA as shown in Fig. 6. This would not be straightforward when using principal components of the whole data space as inputs. By doing so, we have also saved the effort on manually selecting relevant variables that feed into the analysis. We believe that the dimension estimating algorithm we applied minimizes the noise in each of the sub-domains, therefore, achieves the same goal with picking important information manually from each functional domain. Although, the DDR-reduced space would explain less variance than the same dimensional PCA reduced space as PCA is designed to maximize variances; DDR focuses more on the structure within variables that share the same functionality, making sure each functional domain have a representative number of components feeding into CCA.

Both PCA and DDR have their own advantages and disadvantages. Using data that is reduced by PCA as inputs, the canonical correlations are higher than using DDR (Tab. 1 and Tab. 3). Permutation testing gives more significant canonical variables for PCA and those explain slightly higher variance in the original datasets. This is due to the fact that PCA-reduced sub-space is still orthogonal, whereas DDR-reduced sub-space is not. This allows PCA to capture more variance in the observed dataset than DDR (with the same dimensionality). However, DDR saves the effort of selecting relevant variables manually and it automatically estimates the dimensionality. One of the largest drawbacks of PCA is that the results are not as interpretable as DDR. With DDR, we could track the contribution of each sub-domain and directly interpret the canonical loadings of CCA inputs which cannot be easily interpreted in the PCA case. Moreover, these loadings are not subject to the signs of the observed variables. We applied the same stability analysis to canonical loadings on the DDR factors (CCA inputs) and found higher stability than the loadings on the observed variables (Fig. A.17 and Fig. 8).

### What we learn from CCA

In general we have found that with larger sample size, CCA tends to find weaker canonical correlations (Tab. 1 and 3). This is consistent with previous work that showed canonical correlation tends to have higher bias with smaller sample size (Lee [2007]). When we try to interpret canonical correlation using small samples, we should be extremely cautious and depend on out of sample validation to obtain unbiased estimates of canonical correlation.

Additionally, canonical correlations get weaker if we use lower dimensional data as inputs. This is explained by higher dimensional data having greater flexibility to maximize the correlation. We observe that canonical variables constructed by lower dimensional data actually have increased average variance explained in the observed datasets (Tab. 1 and 3). Further analysis shows that the amount of variance explained in the original dataset has a non-linear behavior against CCA input dimension, and it peaks at around dimension 30 in this study.

Further, we found that mean, median and 90th percentile of the distribution of canonical loadings also reduced with increased CCA input dimension. Hence we postulate that higher dimensional inputs may overfit and produce canonical variables that are less related to the original variables.

Since CCA maximizes the correlation between two sets of data rather than the variance canonical variables explain in their original datasets, it is important to be aware that variance explained can be an informative measure, however, cannot become the sole measure used to assess CCA performance. Other measures should be considered such as canonical loadings and canonical correlations.

### Interpretation of CCA loadings

Variable importance is always a major challenge in interpreting CCA results. The canonical weights are the most direct measures of the importance of CCA inputs. However, they are sensitive to the inputs: small perturbation in inputs can lead to significant change in canonical weights, thus not ideal for variable importance evaluation (Bro et al. [2008] and Gittins [2012]). Different studies (Bro et al. [2008] and Thorndike and Weiss [1973]) have suggested using structural coefficients which are also known as canonical loadings to measure the variable importance. In this study, we have shown that it is a stable measurement with the canonical loadings on DDR factors being more stable than on observed variables (Fig. 8 and A.17).

Notably, canonical loadings are sign-subjective since they are just correlations between canonical variables and observed variables/CCA inputs. However, principal components which are often used as the inputs of CCA and canonical variables are sign-invariant, i.e. they may take arbitrary signs between different simulations. Therefore, one should not interpret the absolute sign of canonical loadings as the positive/negative contribution of the variable. We should interpret the loadings as what they are in contrast with, and what is the picture on the other set of canonical variables, in our case, linking SM and BM canonical loadings.

Interpretation of latent factor models like CCA still remains challenging. Researchers often need to trade between model performance and interpretability. However, in the areas of medical/public health research, being able to interpret the results of any statistical/machine learning model is of vital importance. The DDR method proposed here tries to combine prior knowledge on the data collected with mathematical models, to improve the understanding of the intermediate and final results of a CCA pipeline, at the same time, reducing the arbitrary choices researchers have to make and increase analysis automation. To total understand the mechanism between brain and behaviour, and fully interpret the results of all different kinds of latent factor models, much additional research across disciplines is still required.

## Acknowledgments

Zhangdaihong Liu was funded by the Chinese Scholarship Council, EPSRC and MRC (grant number EP/N510129/1); Thomas Nichols was funded by the Wellcome Trust (grant number 100309/Z/12/Z); Stephen Smith was funded by the Wellcome Trust (grant number 110027/Z/15/Z); Kirstie Whitaker was funded by the EPSRC (grant number EP/L015374/1).

## Data Availability Statement

The data that support the findings of this study are available from Human Connectome Project. Restrictions apply to the availability of these data, which were used under license for this study. Data are available at https://db.humanconnectome.org/ with the permission of Human Connectome Project.

## A. Full list of SM variables

The following variables are the 234 subject measures went into the analysis. They are listed using the formal database naming (see https://wiki.humanconnectome.org/display/PublicData/HCP+Data+Dictionary+Public-+Updated+for+the+1200+Subject+Release for their detailed descriptions); names with a ‘-’ sign in front indicates the variable is sign-flipped:

Subject, Release, Acquisition, Gender, Age_in_Yrs, Race_white, Race_black, Race_other, Ethnicity, Height, Weight, BMI, Head_motion, fMRI_3T_ReconVrs, FS_IntraCranial_Vol, FS_BrainSeg_Vol, Handedness, SSAGA_Employ, SSAGA_Income, SSAGA_Educ, SSAGA_InSchool, SSAGA_Rlshp, SSAGA_MOBorn, -SSAGA_BMICat, -SSAGA_BMICatHeaviest, Hematocrit_1, Hematocrit_2, BPSystolic, BPDiastolic, ThyroidHormone, HbA1C, Menstrual_RegCycles, Menstrual_AgeBegan, Menstrual_CycleLength, Menstrual_DaysSinceLast, Menstrual_UsingBirthControl, -FamHist_Moth_Dep, -FamHist_Fath_Dep, -FamHist_Fath_DrgAlc, FamHist_Moth_None, FamHist_Fath_None, -ASR_Anxd_Raw, -ASR_Anxd_Pct, -ASR_Witd_Raw, -ASR_Witd_T, -ASR_Soma_Raw, -ASR_Soma_T, -ASR_Thot_Raw, -ASR_Thot_T, -ASR_Attn_Raw, -ASR_Attn_T, -ASR_Aggr_Raw, -ASR_Aggr_T, -ASR_Rule_Raw, -ASR_Rule_T, -ASR_Intr_Raw, -ASR_Intr_T, -ASR_Oth_Raw, -ASR_Crit_Raw, -ASR_Intn_Raw, -ASR_Intn_T, -ASR_Extn_Raw, -ASR_Extn_T, -ASR_TAO_Sum, -ASR_Totp_Raw, -ASR_Totp_T, -DSM_Depr_Raw, -DSM_Depr_T, -DSM_Anxi_Raw, -DSM_Anxi_T, -DSM_Somp_Raw, -DSM_Somp_T, -DSM_Avoid_Raw, -DSM_Avoid_T, -DSM_Adh_Raw, -DSM_Adh_T, -DSM_Inat_Raw, -DSM_Hype_Raw, -DSM_Antis_Raw, -DSM_Antis_T, -SSAGA_ChildhoodConduct, -SSAGA_PanicDisorder, -SSAGA_Agoraphobia, -SSAGA_Depressive_Ep, -SSAGA_Depressive_Sx, -EVA_Denom, Correction, -Noise_Comp, Odor_Unadj, Odor_AgeAdj, -PainIntens_RawScore, -PainInterf_Tscore, -Taste_Unadj, -Taste_AgeAdj, Mars_Log_Score, -Mars_Errs, Mars_Final, -THC, -SSAGA_Times_Used_Illicits, -SSAGA_Times_Used_Cocaine, -SSAGA_Times_Used_Hallucinogens, -SSAGA_Times_Used_Opiates, -SSAGA_Times_Used_Sedatives, - SSAGA_Times_Used_Stimulants, -SSAGA_Mj_Use, -SSAGA_Mj_Ab_Dep, SSAGA_Mj_Age_1st_Use, - SSAGA_Mj_Times_Used, -Total_Drinks_7days, -Num_Days_Drank_7days, -Avg_Weekday_Drinks_7days, - Avg_Weekend_Drinks_7days, -Total_Beer_Wine_Cooler_7days, -Avg_Weekday_Beer_Wine_Cooler_7days, - Avg_Weekend_Beer_Wine_Cooler_7days, -Total_Wine_7days, -Avg_Weekday Wine_7days, -Avg_Weekend_Wine_7days, -Total_Hard_Liquor_7days, -Avg_Weekday_Hard_Liquor_7days, -Avg_Weekend_Hard_Liquor_7days, -SSAGA_Alc_D4_Dp_Sx, -SSAGA_Alc_D4_Ab_Dx, -SSAGA_Alc_D4_Ab_Sx, -SSAGA_Alc_D4_Dp_Dx, -SSAGA_Alc_12_Drink_Per_Day, SSAGA_Alc_12_Frq, SSAGA_Alc_12_Frq_5plus, SSAGA_Alc_12_Frq_Drk, -SSAGA_Alc_12_Max_Drinks, SSAGA_Alc_Age_1st_Use, -SSAGA_Alc_Hvy_Drinks_Per_Day, SSAGA_Alc_Hvy_Frq, SSAGA_Alc_Hvy_Frq_5plus, SSAGA_Alc_Hvy_Frq_Drk, -SSAGA_Alc_Hvy_Max_Drinks, -Total_Any_Tobacco_7days, -Times_Used_Any_Tobacco_Today, -Num_Days_Used_Any_Tobacco_7days, -Avg_Weekday_Any_Tobacco_7days, -Avg_Weekend_Any_Tobacco_7days, -Total_Cigarettes_7days, -Avg_Weekday_Cigarettes_7days, - Avg_Weekend_Cigarettes_7days, -SSAGA_TB_Smoking_History, -SSAGA_TB_Still_Smoking, MMSE_Score, -PSQI_Score, -PSQI_SleepQuality1, -PSQI_SleepLatency, -PSQI_SleepQuality2, -PSQI_SleepDuration, - PSQI_SleepDisturbance, -PSQI_SleepMeds, -PSQI_DayDysfunction, PicSeq_Unadj, PicSeq_AgeAdj, CardSort_Unadj, CardSort_AgeAdj, Flanker_Unadj, Flanker_AgeAdj, PMAT24_A_CR, -PMAT24_A_SI, -PMAT24_A_RTCR, ReadEng_Unadj, ReadEng_AgeAdj, PicVocab_Unadj, PicVocab_AgeAdj, ProcSpeed_Unadj, ProcSpeed_AgeAdj, DDisc_SV_1mo_200, DDisc_SV_6mo_200, DDisc_SV_1yr_200, DDisc_SV_3yr_200, DDisc_SV_5yr_200, DDisc_SV_10yr_200, DDisc_SV_1mo_40K, DDisc_SV_6mo_40K, DDisc_SV_1yr_40K, DDisc_SV_3yr_40K, DDisc_SV_5yr_40K, DDisc_SV_10yr_40K, DDisc_AUC_200, DDisc_AUC_40K, VSPLOT_TC, -VSPLOT_CRTE, -VSPLOT_OFF, SCPT_TP, SCPT_TN, -SCPT_FP, -SCPT_FN, -SCPT_TPRT, SCPT_SEN, SCPT_SPEC, -SCPT_LRNR, IWRD_TOT, -IWRD_RTC, ListSort_Unadj, ListSort_AgeAdj, ER40_CR, -ER40_CRT, ER40ANG, ER40FEAR, ER40NOE, ER40SAD, -AngAffect_Unadj, -AngHostil_Unadj, -AngAggr_Unadj, -FearAffect_Unadj, -FearSomat_Unadj, -Sadness_Unadj, LifeSatisf_Unadj, MeanPurp_Unadj, PosAffect_Unadj, Friendship_Unadj, -Loneliness_Unadj, -PercHostil_Unadj, -PercReject_Unadj, EmotSupp_Unadj, InstruSupp_Unadj, -PercStress_Unadj, SelfEff_Unadj, Endurance_Unadj, Endurance_AgeAdj, GaitSpeed_Comp, Dexterity_Unadj, Dexterity_AgeAdj, Strength_Unadj, Strength_AgeAdj, NEOFAC_A, NEOFAC_O, NEOFAC_C, NEOFAC_N, NEOFAC_E.

## B. Sign Alignment

### B.1. Effects of Sign-flipping

#### Lemma 1.

*Let X be a column-mean-centred matrix of dimensions N* × *D. Randomly flipping the signs of columns does not affect the subject-wise covariance matrix, and affects the variable-wise covariance matrix in a predictable way*.

*Proof*. Flipping the signs of columns of *X* is equivalent with right-multiply a diagonal matrix *R* with −1 or 1 on diagonal. The subject-wise covariance is defined as<u>^1^</u> 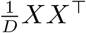. Therefore, flipping the column signs in subject-wise covariance gives

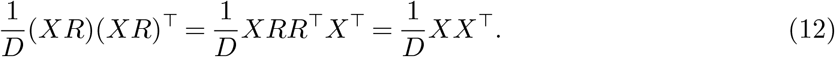

The variable-wise covariance is defined as 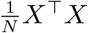. After flipping the column signs,

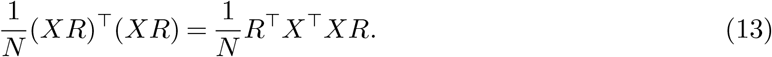

This suggests the variable-wise covariance have corresponding rows and columns sign-flipped simultaneously, i.e. having predictable signs flipped at certain entries. ▀

#### Theorem 2.

*Let X be a column-mean-centred matrix of dimensions N* × *D. Randomly flipping the signs of columns does not change the principal components in PCA, and the principal loadings have the corresponding rows flipped*.

*Proof*. We prove the above statements of PCA by showing the sign-flipping effects on Singular Value Decomposition (SVD). SVD has form

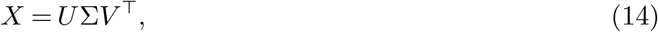

where *U* Σ is known as the principal components and *V* is the principal loadings. Therefore, from the proof of Lemma 1, flipping the column signs of *X* gives

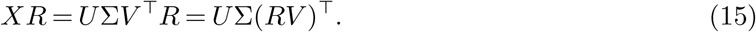

Therefore, it leaves the principal components un-changed and principal loadings have *R* left-multiplied to it which is equivalent with flipping the corresponding row signs. ▀

#### Theorem 3.

*Suppose the inputs of CCA are X and Y*. *Both X and Y are column-mean-centred. Flipping column signs of X and/or Y does not change the canonical variables, but changes the signs of canonical loadings*.

*Proof*. Let *P* and *Q* be canonical variables, *A* and *B* be canonical weights for *X* and *Y* respectively (as described in Section 2.4). We have

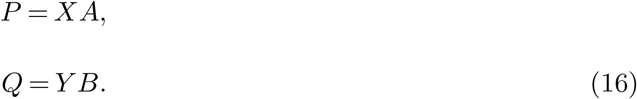

We prove the case for flipping columns of *X* only, and the case for Y can be proved similarly. The solution for *A* are the eigenvectors of 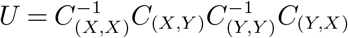, where *C* stands for the covariance matrix. Now we show that flipping column signs of *X* would affect *A* by flipping the respective row signs. We ignore all scaling factors in the covariance matrices. Assume *R* be the sign-flipping diagonal matrix.

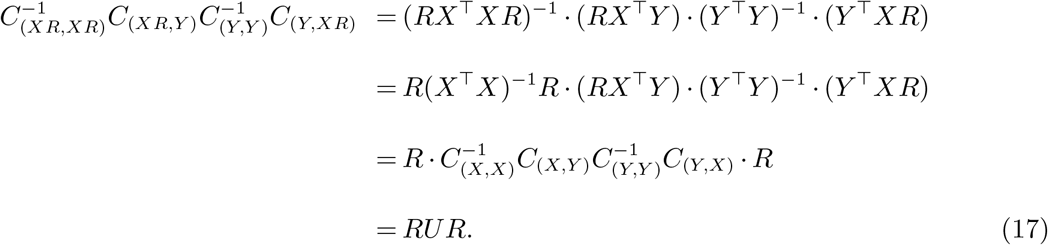

The eigen-decomposition of *RUR* is (ignoring the scaling factor)

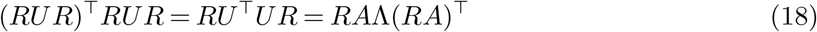

Therefore, by flipping the column signs of *X*, the canonical weights *A* change to *RA*, and the canonical variables *P* remain unchanged (*P* = *XRRA*).

The canonical loadings for *X* are the Pearson’s correlations between *X* and columns of *P*. Thus, by flipping the columns of *X*, we have the canonical loadings sign-flipped for the corresponding variables. ▀

### B.2. Correlation Matrix

**Figure A.1.**
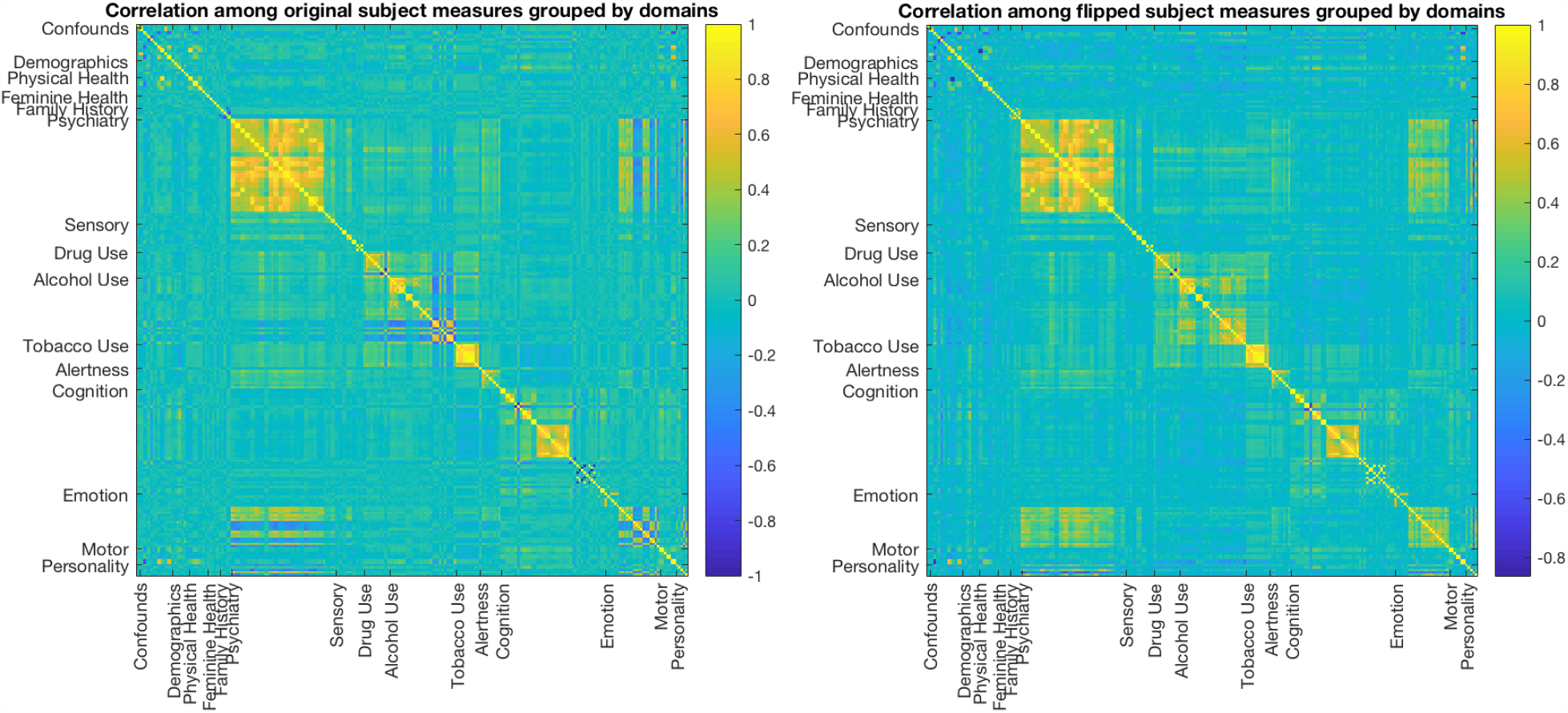
Pairwise correlation among 234 subject measures grouped by 14 functional domains. On the left is the correlation matrix among original variables; on the right is the correlation matrix after sign-flipping which aims to align pairwise correlations between and within domains. Alcohol Use and Emotion sub-domains have the most noticeable changes, reflected by the within domain correlation pattern on the diagonal. However, almost all variables in Tobacco Use and Psychiatry are sign-flipped as well. Flipping all variables within a sub-domain preserves the within-domain correlation pattern. The change is reflected by the correlation with variables in other sub-domains.

## C. Confounders

Confounders include: data release, data acquisition, gender, age (and age^2^), race white (binary), race black (binary), other race (binary), ethnicity, height (and height^2^), weight (and weight^2^), BMI, 3T fMRI Reconstruction Version, head motion, intra-cranial volume (cubed) and brain segmentation volume (cubed).

## D. Canonical loadings for PCA CCA

**Figure A.2.**
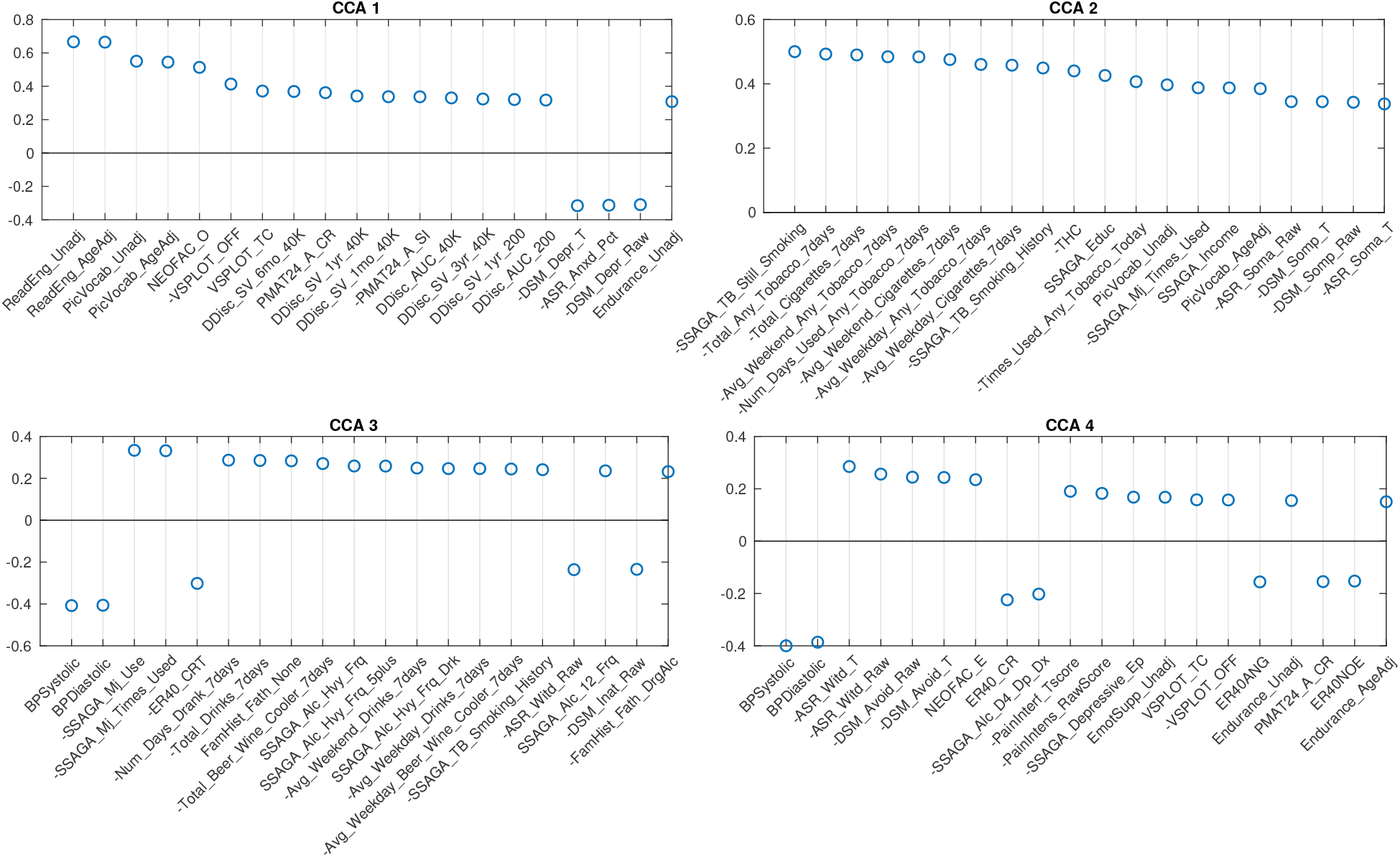
Top 20 canonical loadings for 4 significant SM canonical variables in PCA CCA method using 62 dimensional SM and 100 dimensional BM. Variable names with a ‘-’ sign means the values have been flipped.

## E. Summary Reports

In the following sub-domain reports, by cross-validating PCs within each sub-domain, we observe strong stability on the PCs: PCs in test set explain almost the same amount of variance with the ones in the training set (panel B). This also implies the stability of DDR. We further investigated the principal loadings for the number of DDR estimated dimensions, and found with factor rotation, principal loadings become more interpretable (panel D and E). Therefore, we summarized the meaning of each rotated factor by panel E in the summary reports, and their names are listed in the middle column of Tab. 2.

Moreover, the null eigen-spectrum (green dots in panel A) as introduced in Section 2.3, shows that for example, in the ‘Family History’ sub-domain, only two latent dimensions explain more variance than the ‘background noise’. This is consistent with the DDR estimation for the dimension of ‘Family History’ (panel F, Fig. A.3), and this is the roughly the case for all sub-domains.

**Figure A.3.**
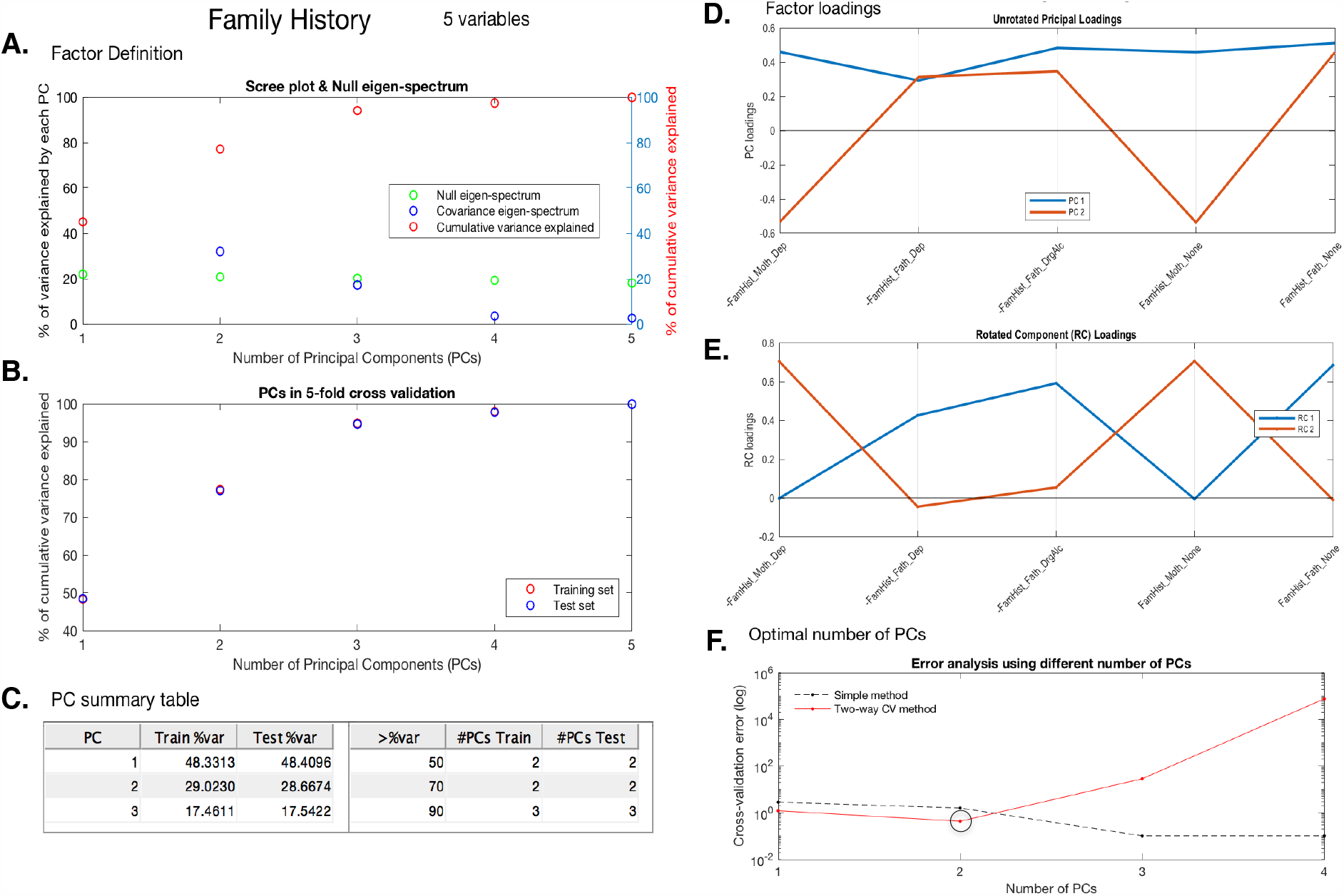
Summary report of Family History. Panel A shows the eigen-spectrum (blue), cumulative eigen-spectrum (red) and null eigen-spectrum (green); panel B shows the cumulative variance explained by principal components (PCs) in cross-validation; panel C is the summary table for panel B showing 3 benchmark percentages 50%, 70% and 90%; panel D shows the principal loadings for optimal number of PCs; panel E shows the rotated loadings in D; panel F shows the error curves calculated by Eqn.6 and Eqn.8, with the minimal error circled at the second component. The naive way of calculating PRESS (dotted line) is monotonically decreasing, while the two-way CV method (red line) offers a minimum point at the second PC (circled).

**Figure A.4.**
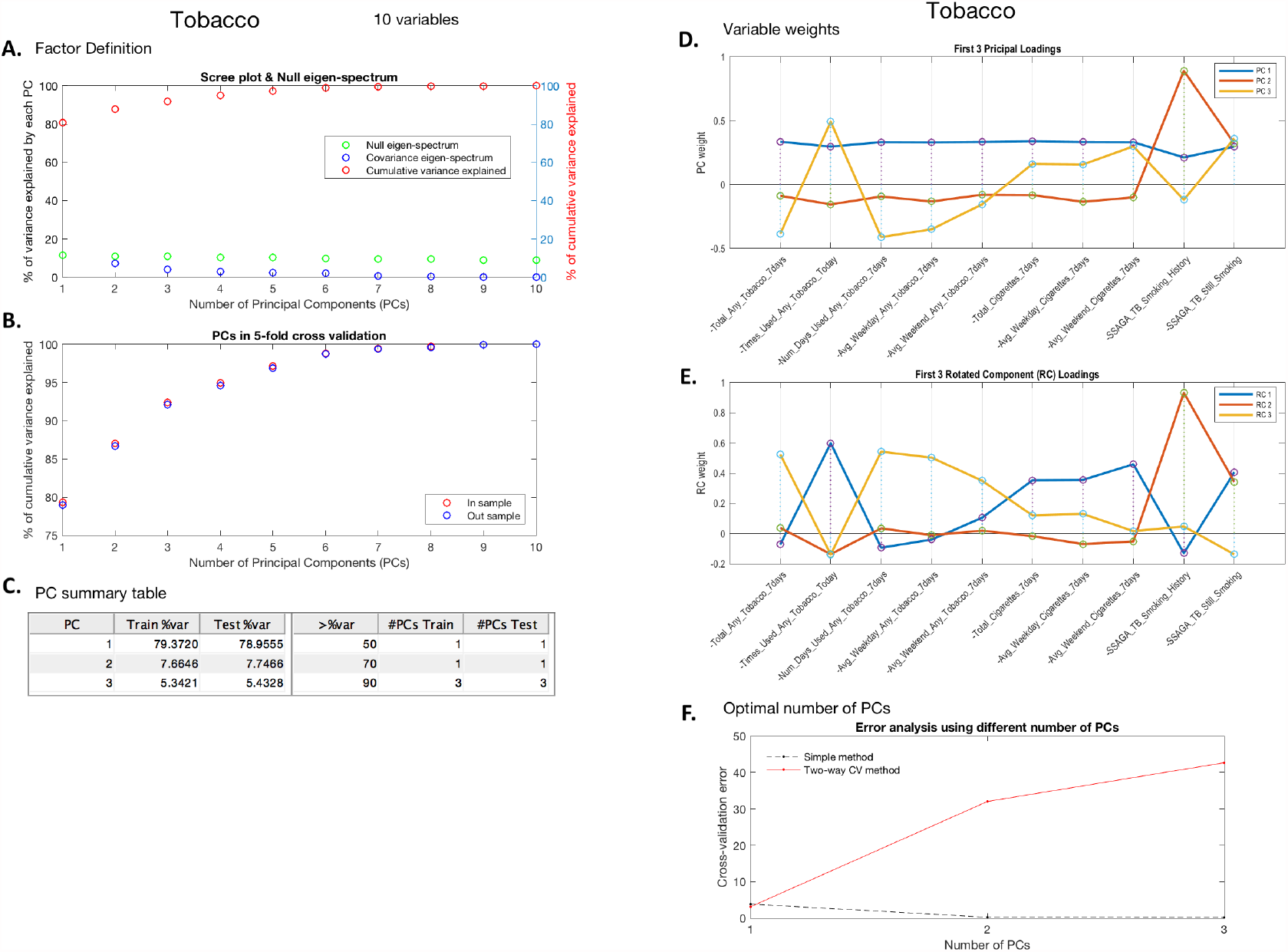
Tobacco Use sub-domain summary report. Panel A shows the eigen-spectrum (blue), cumulative eigen-spectrum (red) and null eigen-spectrum (green); panel B shows the cumulative variance explained by principal components (PCs) in cross-validation; panel C is the summary table for panel B showing 3 benchmark percentages 50%, 70% and 90%; panel D shows the principal loadings for optimal number of PCs; panel E shows the rotated loadings in D; panel F shows the error curves calculated by Eqn.6 and Eqn.8, with the minimal error circled at the second component. The naive way of calculating PRESS (dotted line) is monotonically decreasing, while the two-way CV method (red line) offers a minimum point.

**Figure A.5.**
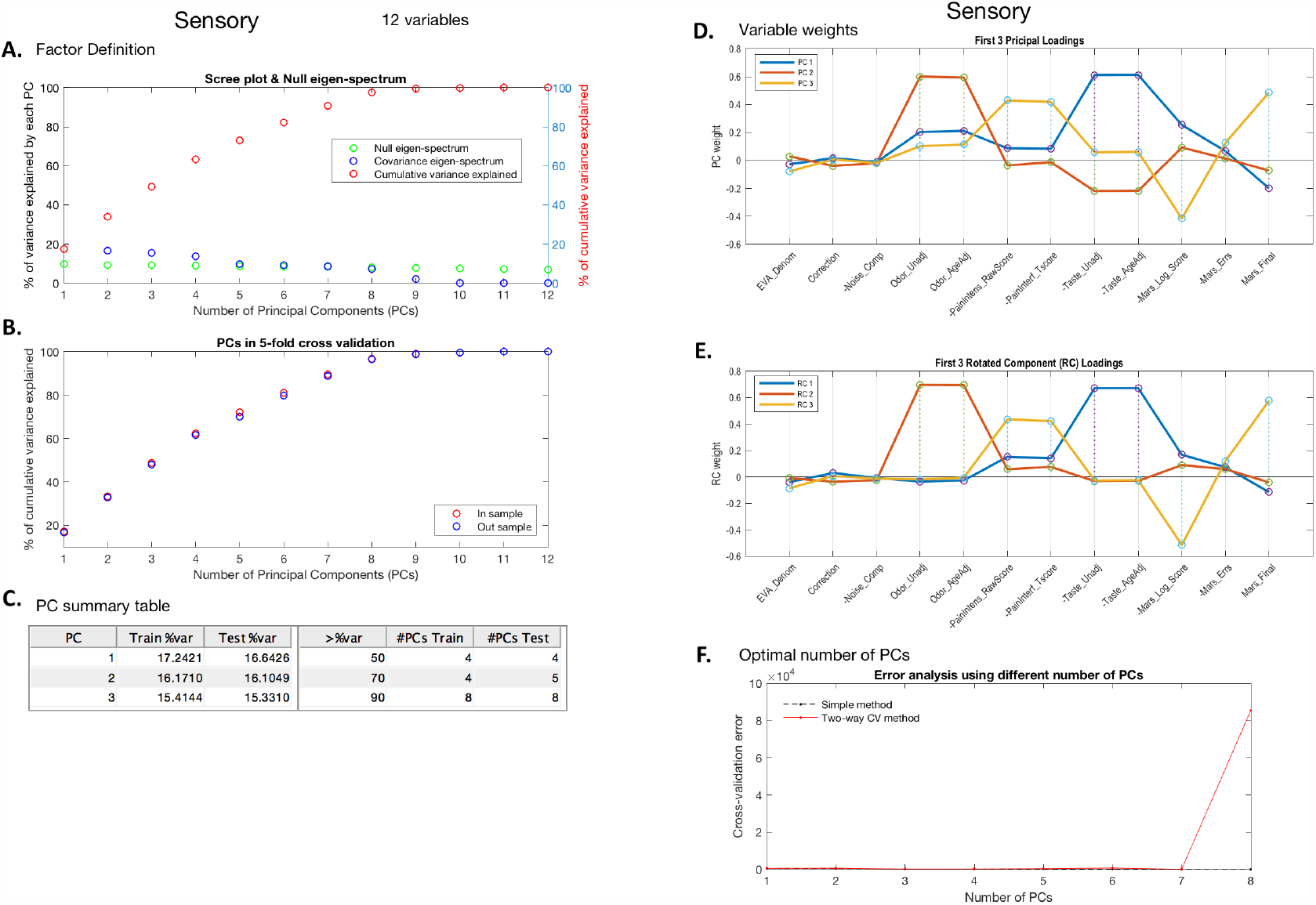
Sensory sub-domain summary report. Panel A shows the eigen-spectrum (blue), cumulative eigen-spectrum (red) and null eigen-spectrum (green); panel B shows the cumulative variance explained by principal components (PCs) in cross-validation; panel C is the summary table for panel B showing 3 benchmark percentages 50%, 70% and 90%; panel D shows the principal loadings for optimal number of PCs; panel E shows the rotated loadings in D; panel F shows the error curves calculated by Eqn.6 and Eqn.8, with the minimal error circled at the second component. The naive way of calculating PRESS (dotted line) is monotonically decreasing, while the two-way CV method (red line) offers a minimum point.

**Figure A.6.**
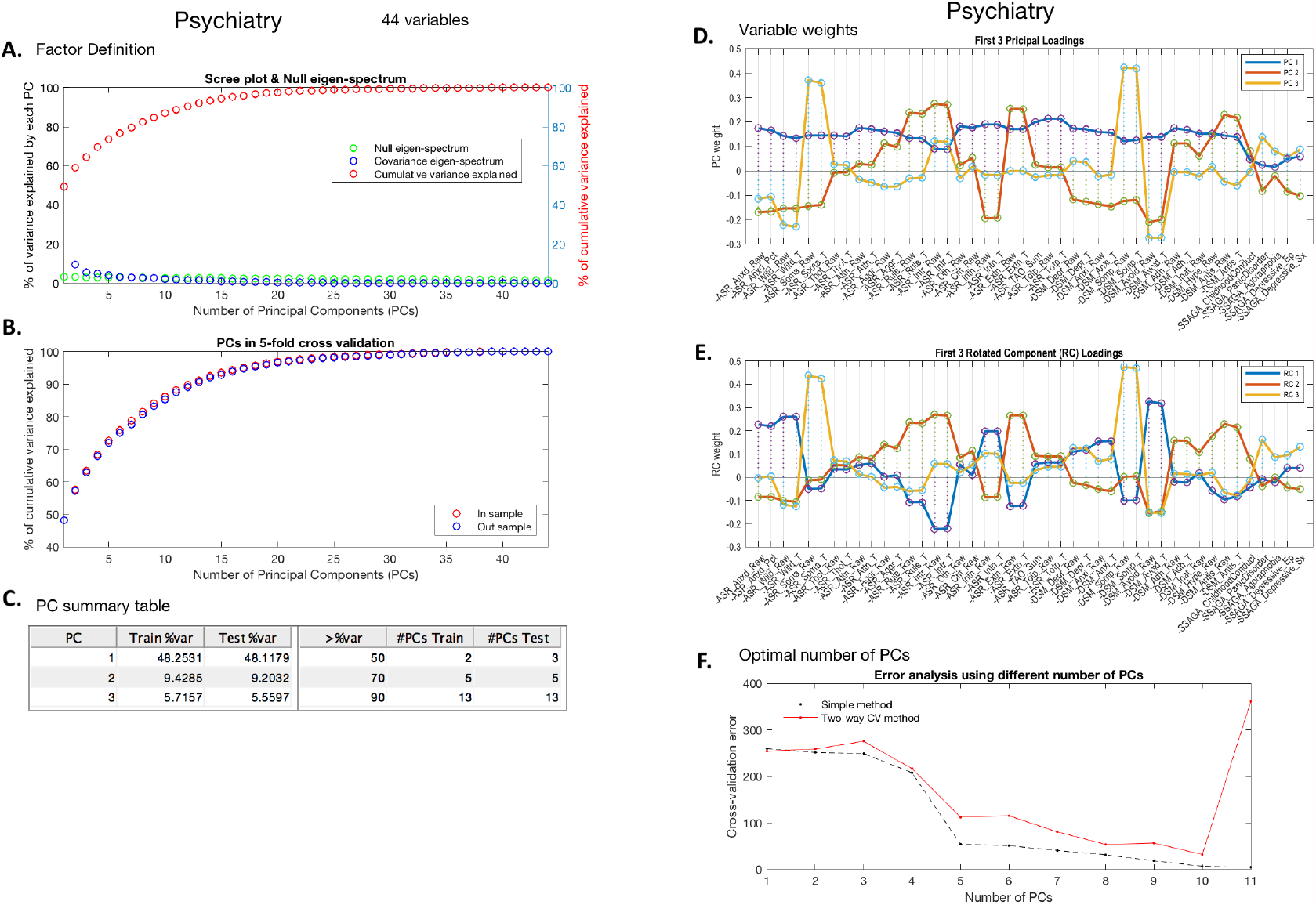
Psychiatry sub-domain summary report. Panel A shows the eigen-spectrum (blue), cumulative eigen-spectrum (red) and null eigen-spectrum (green); panel B shows the cumulative variance explained by principal components (PCs) in cross-validation; panel C is the summary table for panel B showing 3 benchmark percentages 50%, 70% and 90%; panel D shows the principal loadings for optimal number of PCs; panel E shows the rotated loadings in D; panel F shows the error curves calculated by Eqn.6 and Eqn.8, with the minimal error circled at the second component. The naive way of calculating PRESS (dotted line) is monotonically decreasing, while the two-way CV method (red line) offers a minimum point.

**Figure A.7.**
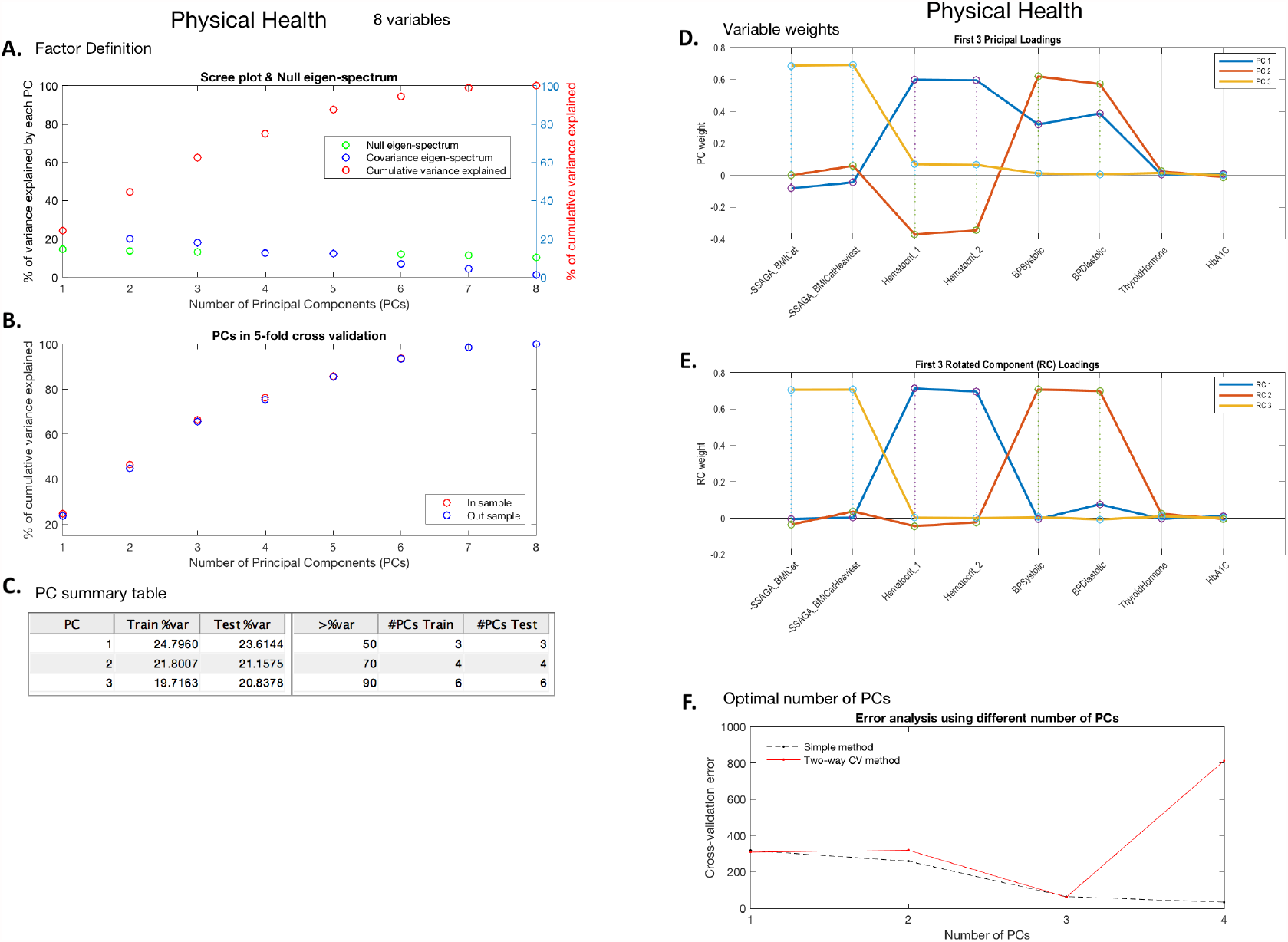
Physical Health sub-domain summary report. Panel A shows the eigen-spectrum (blue), cumulative eigen-spectrum (red) and null eigen-spectrum (green); panel B shows the cumulative variance explained by principal components (PCs) in cross-validation; panel C is the summary table for panel B showing 3 benchmark percentages 50%, 70% and 90%; panel D shows the principal loadings for optimal number of PCs; panel E shows the rotated loadings in D; panel F shows the error curves calculated by Eqn.6 and Eqn.8, with the minimal error circled at the second component. The naive way of calculating PRESS (dotted line) is monotonically decreasing, while the two-way CV method (red line) offers a minimum point.

**Figure A.8.**
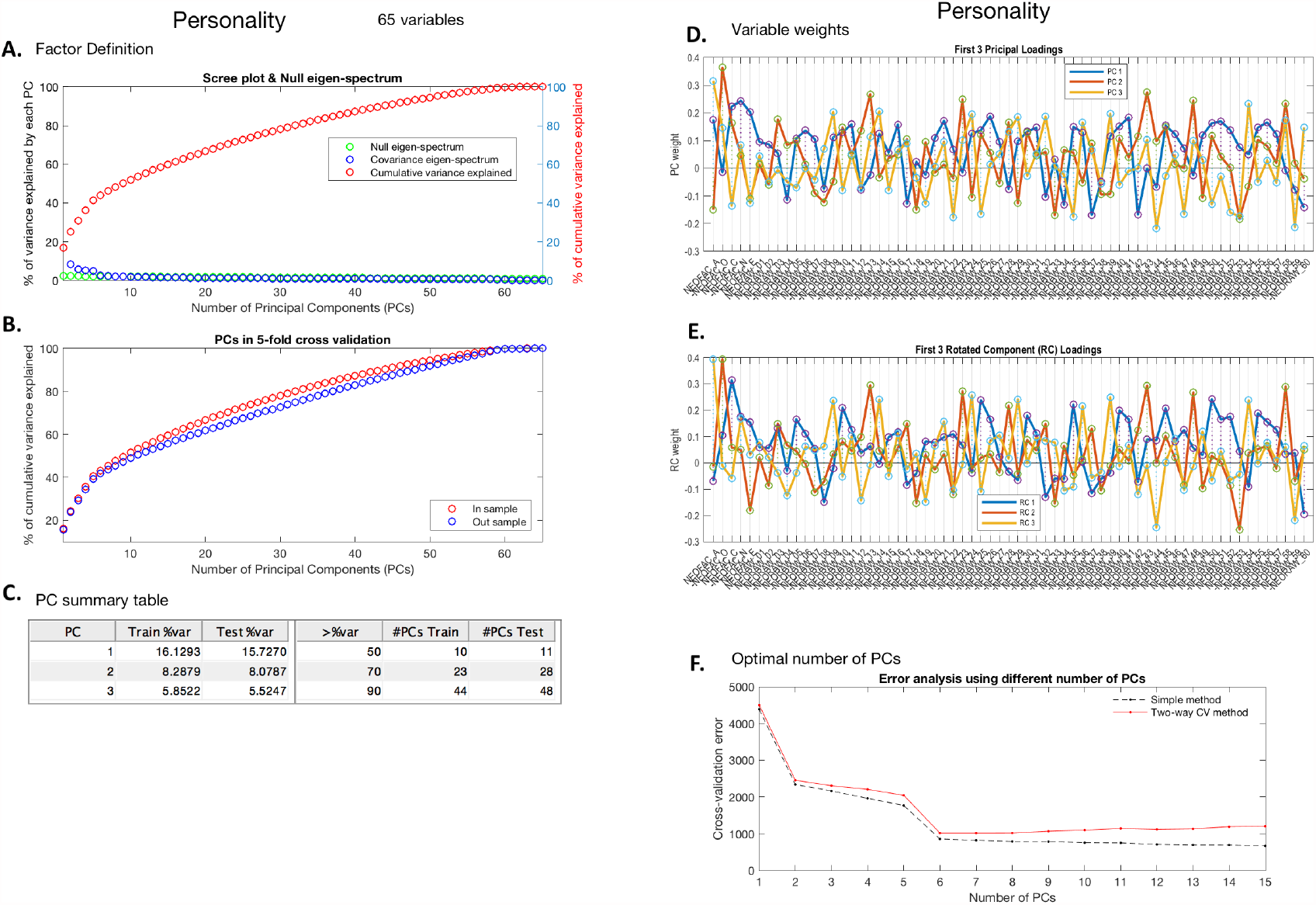
Personality sub-domain summary report. Panel A shows the eigen-spectrum (blue), cumulative eigen-spectrum (red) and null eigen-spectrum (green); panel B shows the cumulative variance explained by principal components (PCs) in cross-validation; panel C is the summary table for panel B showing 3 benchmark percentages 50%, 70% and 90%; panel D shows the principal loadings for optimal number of PCs; panel E shows the rotated loadings in D; panel F shows the error curves calculated by Eqn.6 and Eqn.8, with the minimal error circled at the second component. The naive way of calculating PRESS (dotted line) is monotonically decreasing, while the two-way CV method (red line) offers a minimum point.

**Figure A.9.**
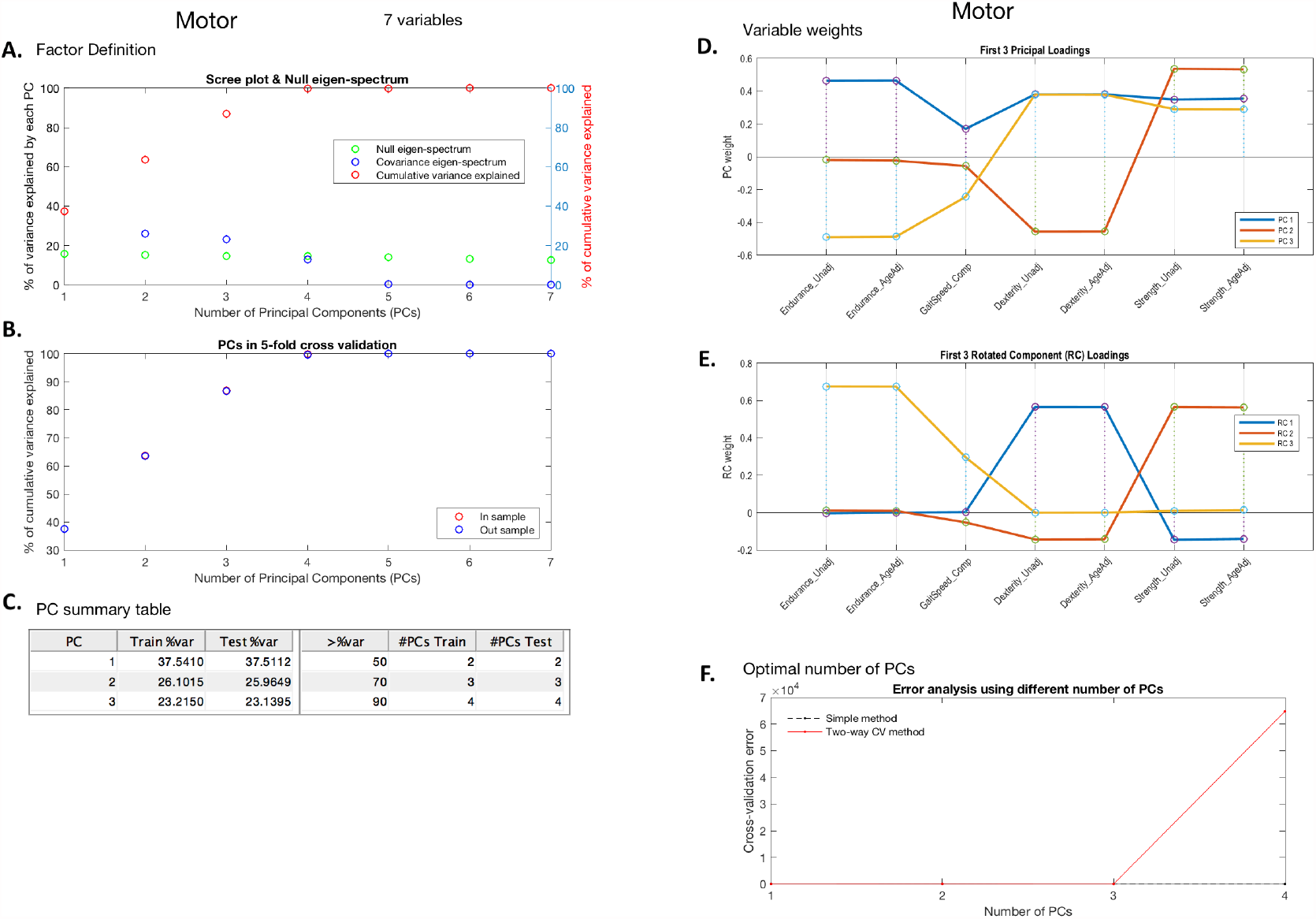
Motor sub-domain summary report. Panel A shows the eigen-spectrum (blue), cumulative eigen-spectrum (red) and null eigen-spectrum (green); panel B shows the cumulative variance explained by principal components (PCs) in cross-validation; panel C is the summary table for panel B showing 3 benchmark percentages 50%, 70% and 90%; panel D shows the principal loadings for optimal number of PCs; panel E shows the rotated loadings in D; panel F shows the error curves calculated by Eqn.6 and Eqn.8, with the minimal error circled at the second component. The naive way of calculating PRESS (dotted line) is monotonically decreasing, while the two-way CV method (red line) offers a minimum point.

**Figure A.10.**
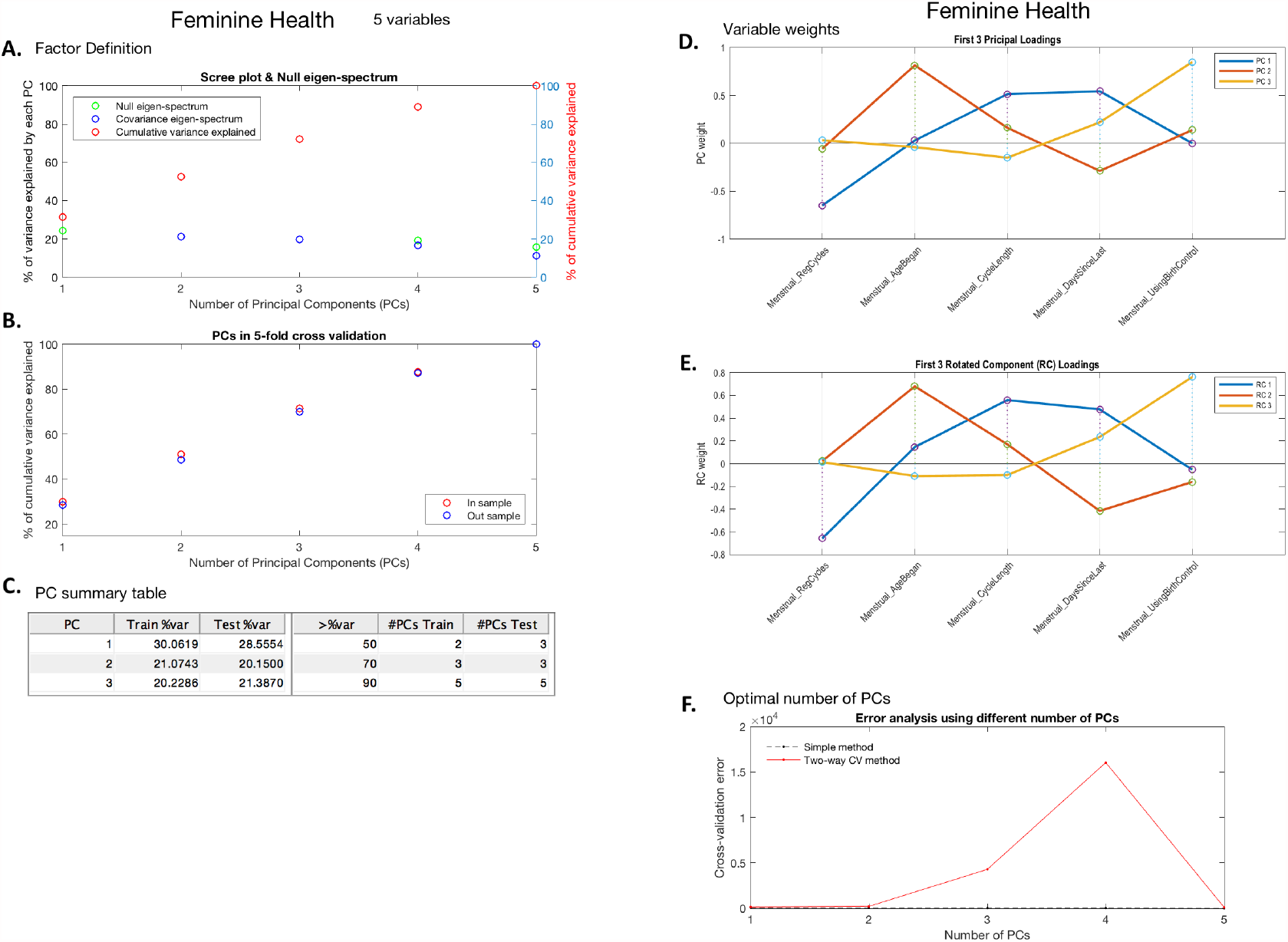
Female Health sub-domain summary report. Notably this sub-domain is generated from female subjects only. Panel A shows the eigen-spectrum (blue), cumulative eigen-spectrum (red) and null eigen-spectrum (green); panel B shows the cumulative variance explained by principal components (PCs) in cross-validation; panel C is the summary table for panel B showing 3 benchmark percentages 50%, 70% and 90%; panel D shows the principal loadings for optimal number of PCs; panel E shows the rotated loadings in D; panel F shows the error curves calculated by Eqn.6 and Eqn.8, with the minimal error circled at the second component. The naive way of calculating PRESS (dotted line) is monotonically decreasing, while the two-way CV method (red line) offers a minimum point.

**Figure A.11.**
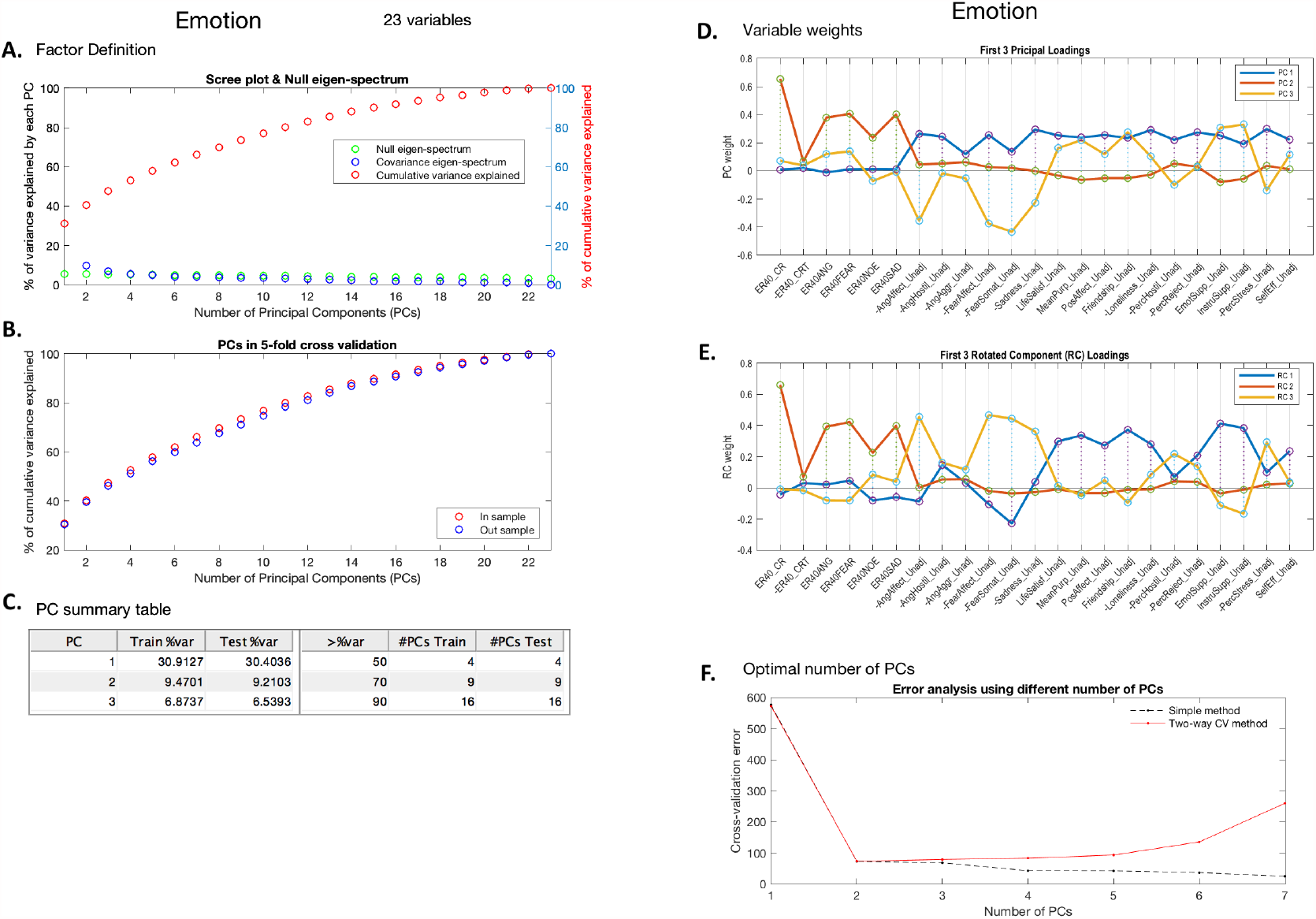
Emotion sub-domain summary report. Panel A shows the eigen-spectrum (blue), cumulative eigen-spectrum (red) and null eigen-spectrum (green); panel B shows the cumulative variance explained by principal components (PCs) in cross-validation; panel C is the summary table for panel B showing 3 benchmark percentages 50%, 70% and 90%; panel D shows the principal loadings for optimal number of PCs; panel E shows the rotated loadings in D; panel F shows the error curves calculated by Eqn.6 and Eqn.8, with the minimal error circled at the second component. The naive way of calculating PRESS (dotted line) is monotonically decreasing, while the two-way CV method (red line) offers a minimum point.

**Figure A.12.**
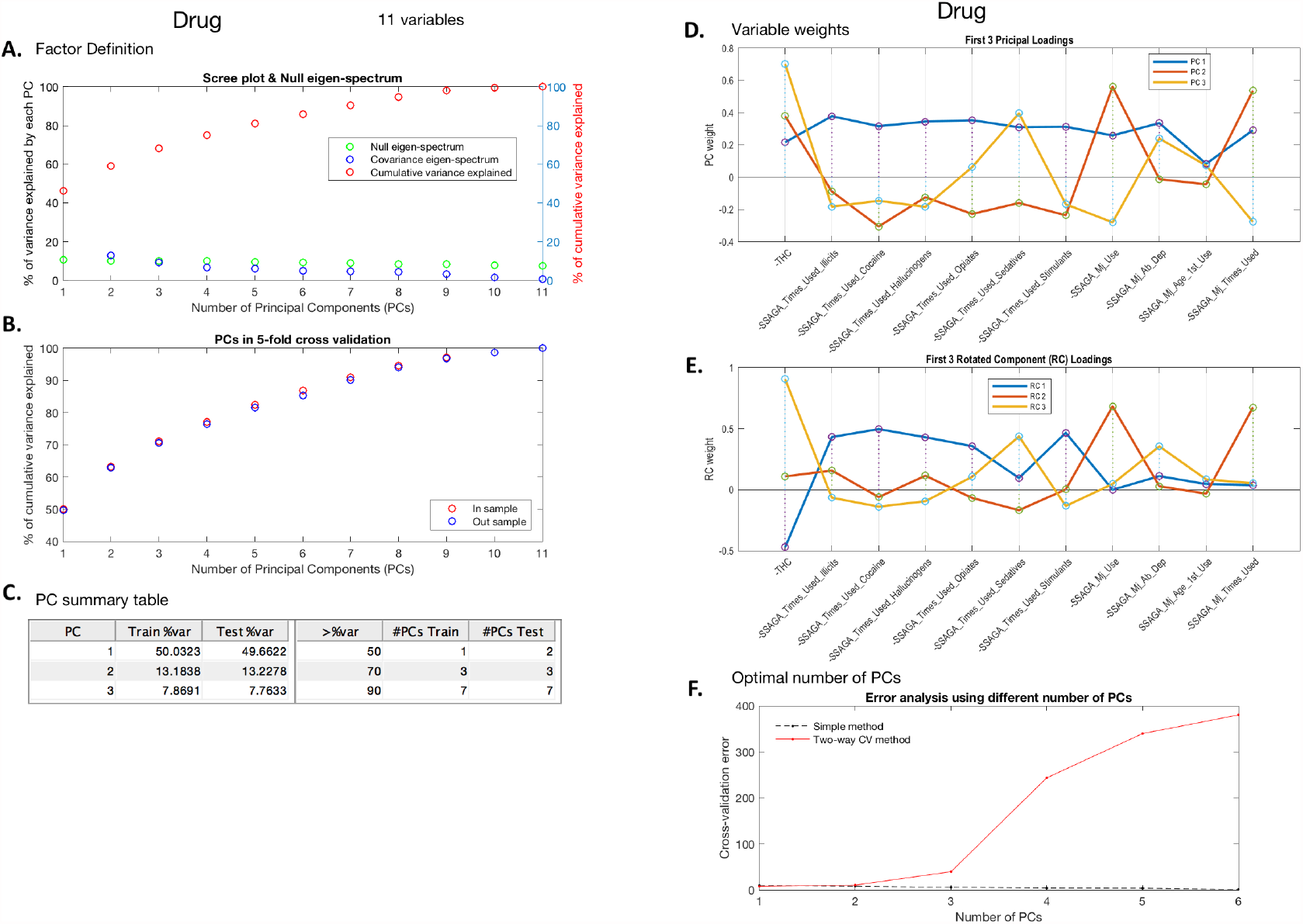
Drug Use sub-domain summary report. Panel A shows the eigen-spectrum (blue), cumulative eigen-spectrum (red) and null eigen-spectrum (green); panel B shows the cumulative variance explained by principal components (PCs) in cross-validation; panel C is the summary table for panel B showing 3 benchmark percentages 50%, 70% and 90%; panel D shows the principal loadings for optimal number of PCs; panel E shows the rotated loadings in D; panel F shows the error curves calculated by Eqn.6 and Eqn.8, with the minimal error circled at the second component. The naive way of calculating PRESS (dotted line) is monotonically decreasing, while the two-way CV method (red line) offers a minimum point.

**Figure A.13.**
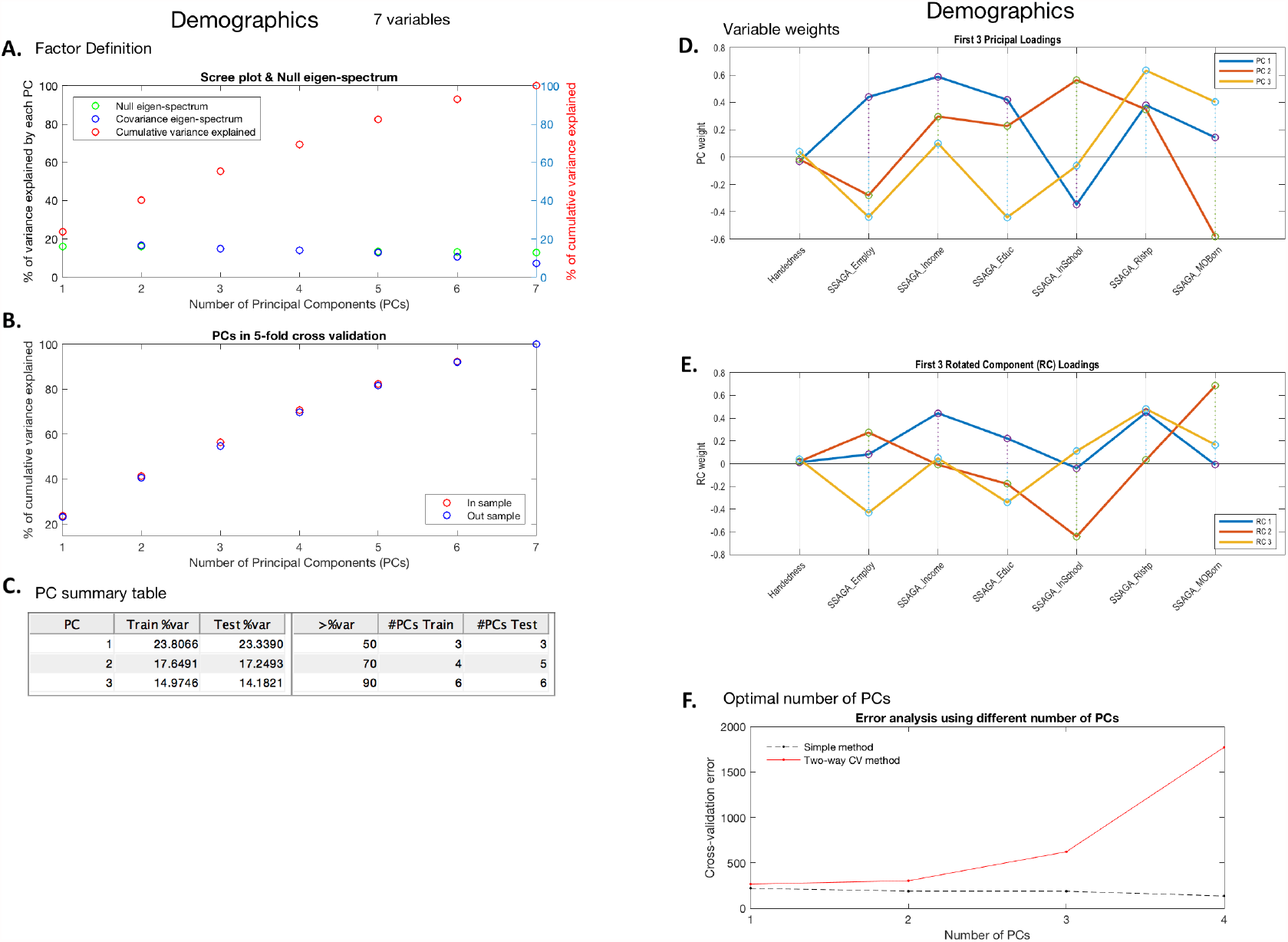
Demographics and SES sub-domain summary report. Panel A shows the eigen-spectrum (blue), cumulative eigen-spectrum (red) and null eigen-spectrum (green); panel B shows the cumulative variance explained by principal components (PCs) in cross-validation; panel C is the summary table for panel B showing 3 benchmark percentages 50%, 70% and 90%; panel D shows the principal loadings for optimal number of PCs; panel E shows the rotated loadings in D; panel F shows the error curves calculated by Eqn.6 and Eqn.8, with the minimal error circled at the second component. The naive way of calculating PRESS (dotted line) is monotonically decreasing, while the two-way CV method (red line) offers a minimum point.

**Figure A.14.**
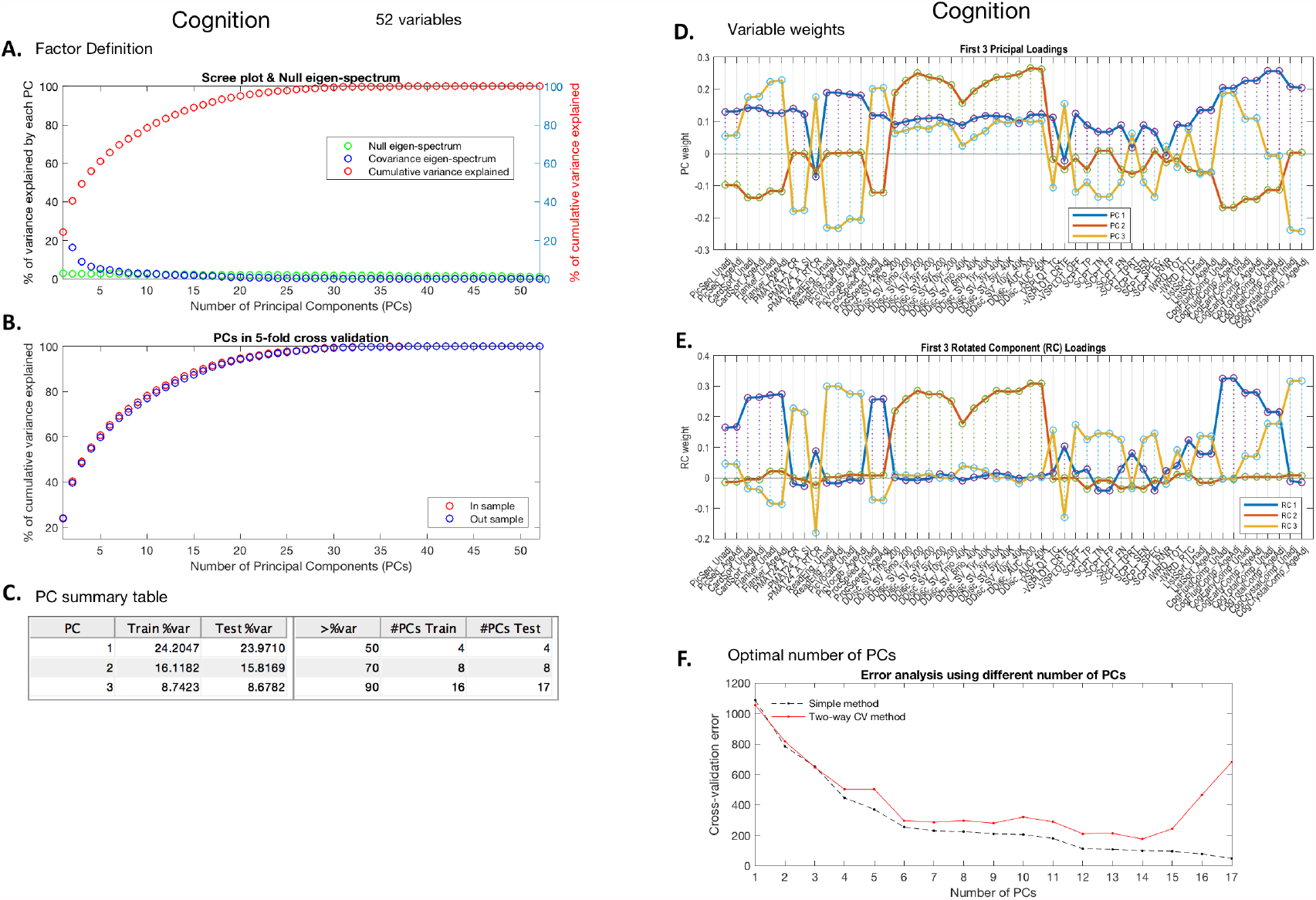
Cognition sub-domain summary report. Panel A shows the eigen-spectrum (blue), cumulative eigen-spectrum (red) and null eigen-spectrum (green); panel B shows the cumulative variance explained by principal components (PCs) in cross-validation; panel C is the summary table for panel B showing 3 benchmark percentages 50%, 70% and 90%; panel D shows the principal loadings for optimal number of PCs; panel E shows the rotated loadings in D; panel F shows the error curves calculated by Eqn.6 and Eqn.8, with the minimal error circled at the second component. The naive way of calculating PRESS (dotted line) is monotonically decreasing, while the two-way CV method (red line) offers a minimum point.

**Figure A.15.**
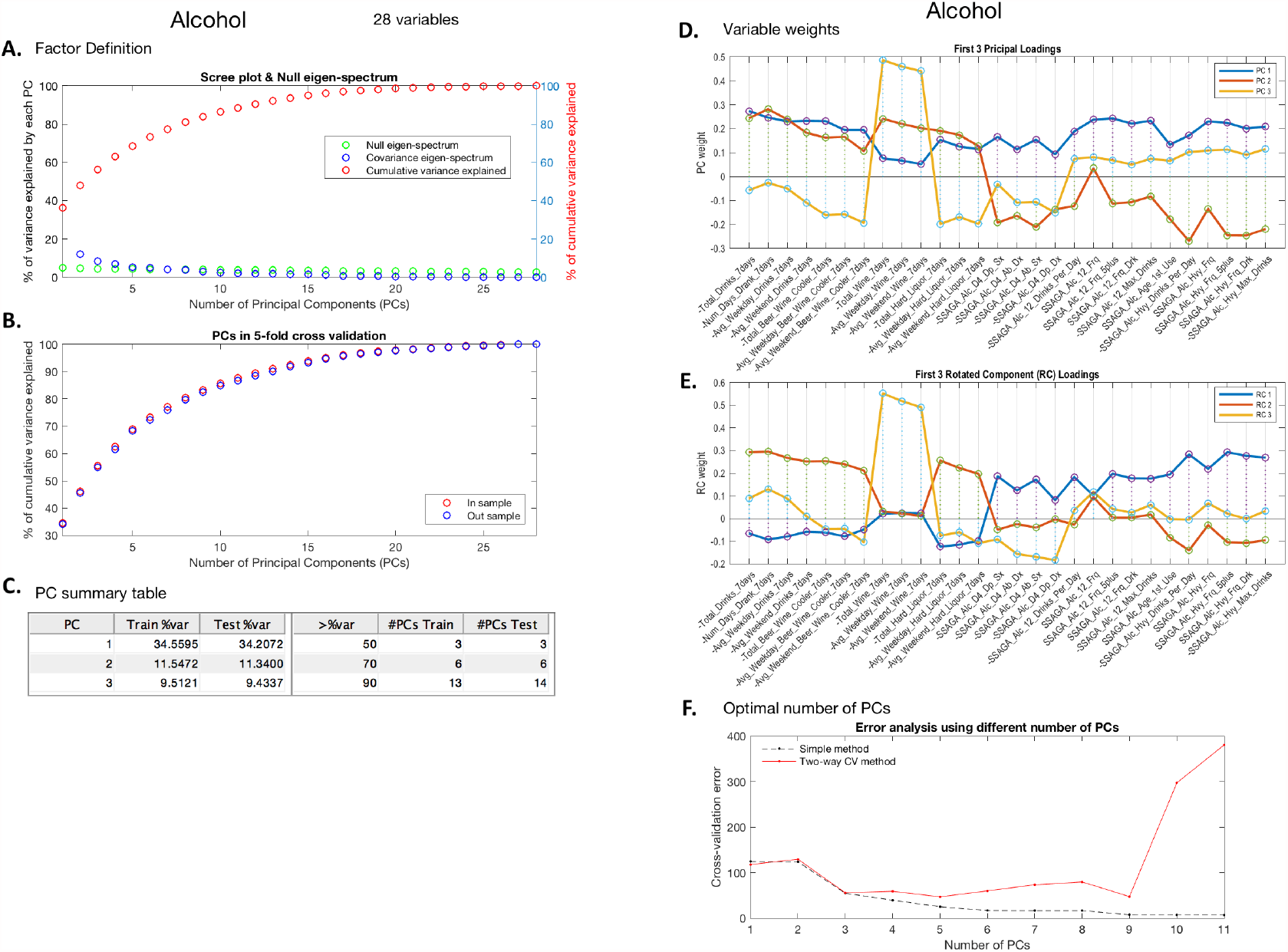
Alcohol Use sub-domain summary report. Panel A shows the eigen-spectrum (blue), cumulative eigen-spectrum (red) and null eigen-spectrum (green); panel B shows the cumulative variance explained by principal components (PCs) in cross-validation; panel C is the summary table for panel B showing 3 benchmark percentages 50%, 70% and 90%; panel D shows the principal loadings for optimal number of PCs; panel E shows the rotated loadings in D; panel F shows the error curves calculated by Eqn.6 and Eqn.8, with the minimal error circled at the second component. The naive way of calculating PRESS (dotted line) is monotonically decreasing, while the two-way CV method (red line) offers a minimum point.

**Figure A.16.**
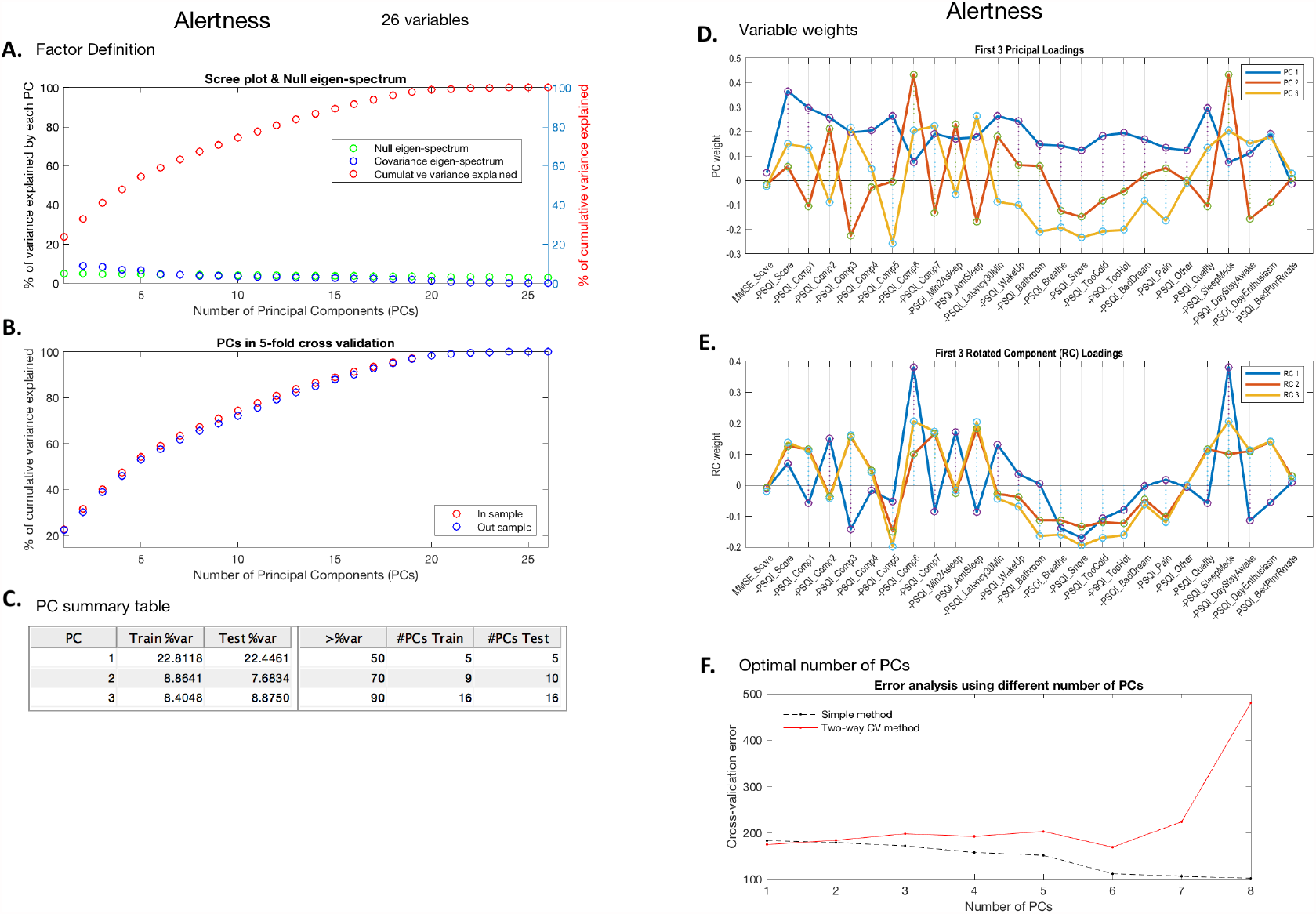
Alertness sub-domain summary report. Panel A shows the eigen-spectrum (blue), cumulative eigen-spectrum (red) and null eigen-spectrum (green); panel B shows the cumulative variance explained by principal components (PCs) in cross-validation; panel C is the summary table for panel B showing 3 benchmark percentages 50%, 70% and 90%; panel D shows the principal loadings for optimal number of PCs; panel E shows the rotated loadings in D; panel F shows the error curves calculated by Eqn.6 and Eqn.8, with the minimal error circled at the second component. The naive way of calculating PRESS (dotted line) is monotonically decreasing, while the two-way CV method (red line) offers a minimum point.

## F. Stability of canonical loadings

**Figure A.17.**
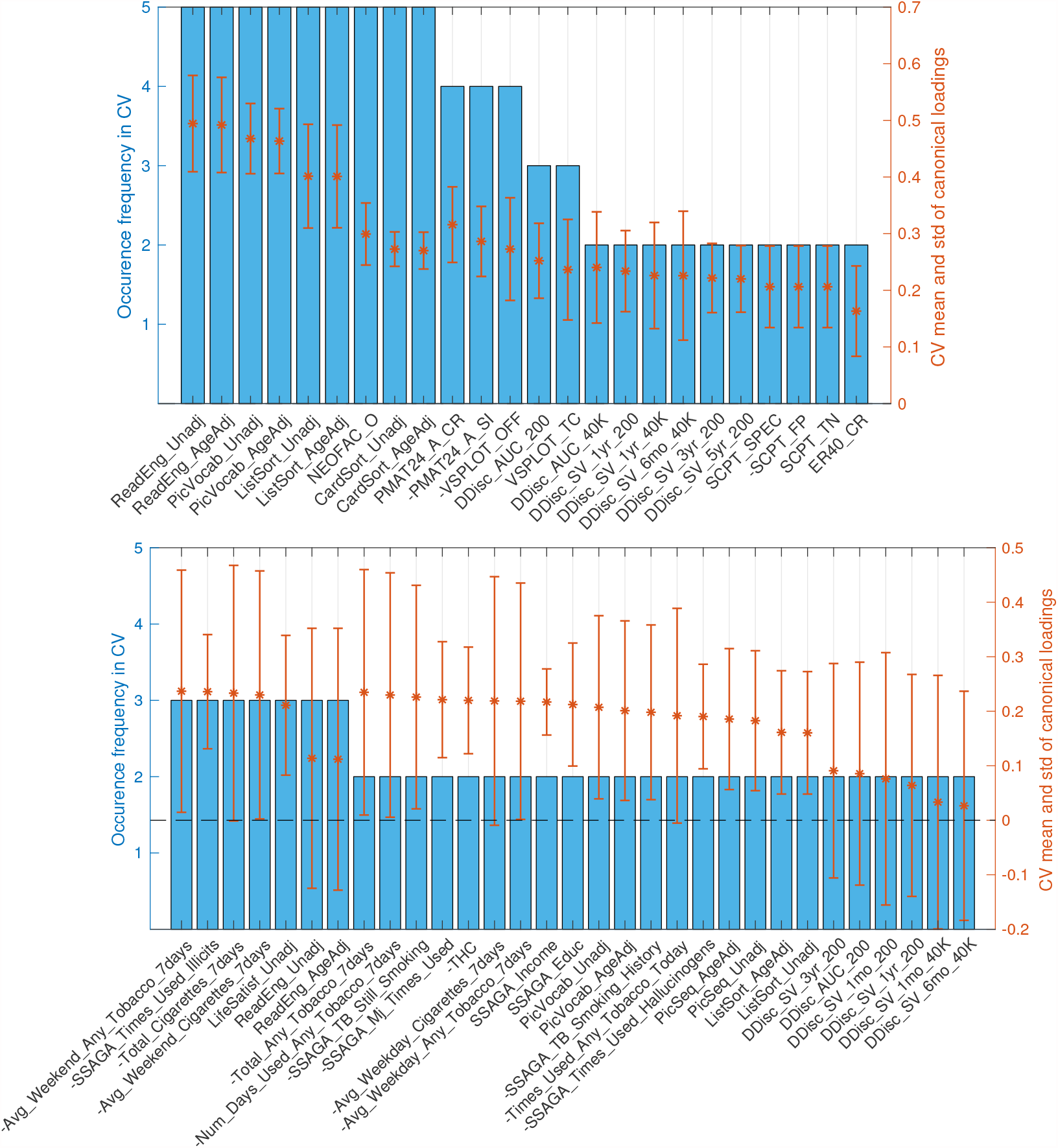
Stability of SM canonical loadings on observed variables. Bar plot shows the occurrence frequency in CV out of the 5 folds. Variables are chosen by selecting the top 20 mostly weighted ones in each fold. The ones appeared at least twice are shown above. Right axis shows the mean and the standard deviation over all occurred loadings. Top and bottom plots are the canonical loadings for the first and second canonical variables respectively. It is obvious that the second canonical loadings are less stable than the first set.

**Figure A.18.**
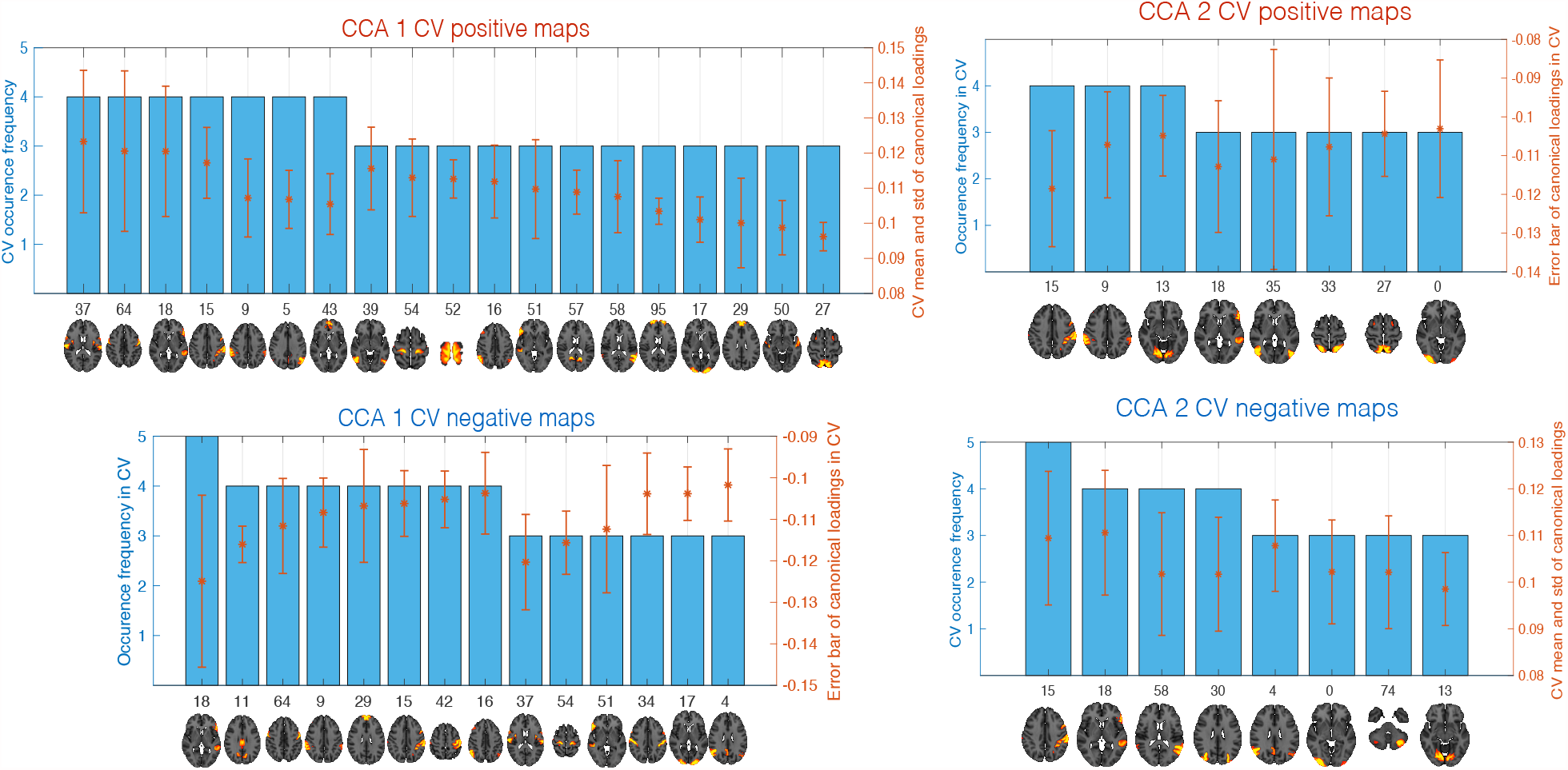
Stability of BM canonical loadings on observed data. Bar plot shows the occurrence frequency in CV out of the 5 folds. The positive (top plots) and negative (bottom plots) maps are chosen by first averaging the top 20 positive and negative canonical loadings within each region respectively; then select the top 20 nodes with the highest positive and negative mean loadings in each fold. The ones occurred at least three times are shown above. Right axis shows the mean and the standard deviation over all occurred loadings. Similar to SM canonical loadings, the first set shows better stability than the second set.

## Footnotes

1 Positive map: http://neurosynth.org/decode/?neurovault=108956; negative map: http://neurosynth.org/decode/?neurovault=108957

2 Positive map: http://neurosynth.org/decode/?neurovault=108976; negative map: http://neurosynth.org/decode/?neurovault=108977

3 Positive map: http://neurosynth.org/decode/?neurovault=108978; negative map: http://neurosynth.org/decode/?neurovault=108979

## References

David C Van Essen, Stephen M Smith, Deanna M Barch, Timothy EJ Behrens, Essa Yacoub, Kamil Ugurbil, Wu-Minn HCP Consortium, et al. The wu-minn human connectome project: an overview. Neuroimage, 80:62–79, 2013.

Cathie Sudlow, John Gallacher, Naomi Allen, Valerie Beral, Paul Burton, John Danesh, Paul Downey, Paul Elliott, Jane Green, Martin Landray, et al. Uk biobank: an open access resource for identifying the causes of a wide range of complex diseases of middle and old age. PLoS medicine, 12(3):e1001779, 2015.

Harold Hotelling. Relations between two sets of variates. Biometrika, 28(3/4):321–377, 1936.

Bruce Thompson. Canonical correlation analysis. Encyclopedia of statistics in behavioral science, 2005.

Stephen M Smith, Thomas E Nichols, Diego Vidaurre, Anderson M Winkler, Timothy EJ Behrens, Matthew F Glasser, Kamil Ugurbil, Deanna M Barch, David C Van Essen, and Karla L Miller. A positive-negative mode of population covariation links brain connectivity, demographics and behavior. Nature neuroscience, 18(11):1565–1567, 2015.

Kuldeep Kumar, Laurent Chauvin, Matthew Toews, Olivier Colliot, and Christian Desrosiers. Multi-modal brain fingerprinting: a manifold approximation based framework. bioRxiv, page 209726, 2017.

Anjali Krishnan, Lynne J Williams, Anthony Randal McIntosh, and Hervé Abdi. Partial least squares (pls) methods for neuroimaging: a tutorial and review. Neuroimage, 56(2):455–475, 2011.

Jing Sui, Tülay Adali, Godfrey Pearlson, Honghui Yang, Scott R Sponheim, Tonya White, and Vince D Calhoun. A cca+ ica based model for multi-task brain imaging data fusion and its application to schizophrenia. Neuroimage, 51(1):123–134, 2010.

Diego Vidaurre, Stephen M Smith, and Mark W Woolrich. Brain network dynamics are hierarchically organized in time. Proceedings of the National Academy of Sciences, 114(48):12827–12832, 2017.

Ola Friman, Jonny Cedefamn, Peter Lundberg, Magnus Borga, and Hans Knutsson. Detection of neural activity in functional mri using canonical correlation analysis. Magnetic Resonance in Medicine, 45(2):323–330, 2001.

Claudia Grellmann, Sebastian Bitzer, Jane Neumann, Lars T Westlye, Ole A Andreassen, Arno Villringer, and Annette Horstmann. Comparison of variants of canonical correlation analysis and partial least squares for combined analysis of mri and genetic data. Neuroimage, 107:289–310, 2015.

Kirstie J Whitaker, Petra E Vértes, Rafael Romero-Garcia, František Váša, Michael Moutoussis, Gita Prabhu, Nikolaus Weiskopf, Martina F Callaghan, Konrad Wagstyl, Timothy Rittman, et al. Adolescence is associated with genomically patterned consolidation of the hubs of the human brain connectome. Proceedings of the National Academy of Sciences, 113(32):9105–9110, 2016.

Stephen M Smith, Christian F Beckmann, Jesper Andersson, Edward J Auerbach, Janine Bijsterbosch, Gwenäelle Douaud, Eugene Duff, David A Feinberg, Ludovica Griffanti, Michael P Harms, et al. Resting-state fmri in the human connectome project. Neuroimage, 80:144–168, 2013.

Andrei Nikolaevich Tikhonov. On the solution of ill-posed problems and the method of regularization. In Doklady Akademii Nauk, volume 151, pages 501–504. Russian Academy of Sciences, 1963.

G Blom. Statistical elements and transformed beta variables. Wiley, New York.

Boeschen, LE, Koss, MP, Figueredo, AJ, & Coan, JA (2001). Experiential avoidance and post-traumatic stress disorder: A cognitive mediational model of rape recovery. Journal of Aggression, Maltreatment & Trauma, 4(2):211–245, 1958.

T Mark Beasley, Stephen Erickson, and David B Allison. Rank-based inverse normal transformations are increasingly used, but are they merited? Behavior genetics, 39(5):580, 2009.

R. Bro, K. Kjeldahl, A. K. Smilde, and H. A. L. Kiers. Cross-validation of component models: A critical look at current methods. Analytical and Bioanalytical Chemistry, 390(5):1241–1251, 2008. ISSN 1618-2642. doi: 10.1007/s00216-007-1790-1. URL http://link.springer.com/10.1007/s00216-007-1790-1.

Magnus Borga. Canonical correlation: a tutorial. On line tutorial http://people.imt.liu.se/magnus/cca, pages 1–12, 2001. ISSN 1940-6029. doi: 10.1007/978-1-60761-977-26. URL http://people.imt.liu.se/{~}magnus/cca/tutorial/tutorial.pdf.

Boris Egloff, Andreas Schwerdtfeger, Stefan C Schmukle, and Johannes Gutenberg-university Mainz. Conducting and Interpreting Canonical Correlation Analysis in Personality Research: A uSE. Journal of Personality Assessment, 84(1):82–88, 2010. ISSN 0022-3891. doi: 10.1207/s15327752jpa8401.

Henry F Kaiser. The varimax criterion for analytic rotation in factor analysis. Psychometrika, 23 (3):187–200, 1958.

HS Lee. Canonical correlation analysis using small number of samples. Communications in Statistics-Simulation and Computation, 36(5):973–985, 2007. ISSN 0361-0918. doi: 10.1080/03610910701539443.

Robert Gittins. Canonical analysis: a review with applications in ecology, volume 12. Springer Science & Business Media, 2012.

Robert M Thorndike and David J Weiss. A study of the stability of canonical correlations and canonical components. Educational and Psychological Measurement, 33(1):123–134, 1973.

